# STMN2 protein depletion via translation deficits and stress granules and its compensation in ALS

**DOI:** 10.64898/2026.01.25.701321

**Authors:** Brittany C. S. Ellis, Anna Sanchez Avila, Wan-Ping Huang, Sabin J. John, Sam Bonsall, Rachel E. Hodgson, Vedanth Kumar, Matthew Nolan, Susan G. Campbell, Kurt J. De Vos, Clotilde Lagier-Tourenne, J. Robin Highley, Johnathan Cooper-Knock, Tatyana A. Shelkovnikova

## Abstract

STMN2 is an abundant neurospecific protein dysregulated in neurodegenerative diseases such as amyotrophic lateral sclerosis (ALS). We previously reported that cellular stress can lead to STMN2 loss due to TDP-43 nuclear condensation. Here, using human and murine neuronal cell models, multiple pharmacological tools, single-molecule *in situ* analysis of mRNA localisation and translation, and longitudinal analysis of neuronal fitness/survival, we establish TDP-43-independent mechanisms of STMN2 depletion linked to stress response. We find that human STMN2 protein is extremely labile under acute high-magnitude stress. Early in stress, STMN2 is suppressed via activated proteasomal degradation, phosphorylation and translational repression by stress granules, independently of TDP-43 loss of function in splicing. We further show that STMN2 protein level is highly sensitive to chronic translation deficits, such as those elicited by prolonged low-grade stress. Finally, we demonstrate that STMN2 mRNA is upregulated in non-TDP ALS such as ALS-FUS, which may compensate for translation/stress granule defects in these disease subtypes. Consistent with the compensation hypothesis, STMN2 mRNA is also upregulated in the relatively spared (cortex), but not severely affected (spinal cord), CNS regions in ALS-TDP. In conclusion, our study implicates two common hallmarks of neurodegeneration, translation impairment and abnormal stress granules, in STMN2 depletion and reports an RNA-level compensation that fails in neurons with TDP-43 pathology. Our study supports the development of stress response targeting therapies in ALS with and without TDP-43 pathology.

## INTRODUCTION

A neurospecific protein STMN2 has recently emerged as a molecular factor dysregulated in neurodegenerative conditions with TDP-43 proteinopathy, including subsets of amyotrophic lateral sclerosis (ALS), frontotemporal dementia (FTD), and Alzheimer’s disease (1–3). Mechanistically, STMN2 pre-mRNA has TDP-43 binding sites in the vicinity of a cryptic exon (CE; exon 2a) in intron 1, and this CE is preferentially spliced in upon TDP-43 dysregulation. This results in STMN2 mRNA truncation, ultimately leading to reduced STMN2 protein synthesis (4). STMN2 has also been implicated in ALS with SOD1 mutations, spinal muscular atrophy (5) and Parkinson’s disease (6). Stmn2 knockout causes neuropathy in mice (7–9). Thus, loss of STMN2 homeostasis may act as a convergent pathway of neuronal degeneration across several neurological diseases.

STMN2 (aka SCG10) is a conserved phosphoprotein (10) expressed from one of the 25 most abundant mRNAs in human and rodent motor neurons (11). It has an important role in neurite outgrowth, being highly expressed in the developing neurons (12), and is recruited to the growth cones of regenerating axons after axonal injury (13). STMN2 inhibits microtubule polymerisation and therefore ensures the dynamism of this process – required for both neurite growth in the development and axonal regeneration in the adult state (14).

We have recently demonstrated that acute cellular stress, such as short exposure to arsenite, results in rapid and profound STMN2 downregulation, correlating with TDP-43 nuclear condensation and loss of solubility (15). Importantly, although stress-induced changes in TDP-43-regulated splicing impact multiple TDP-43 targets, STMN2 is particularly sensitive (15). Stress may play an important role in neurodegenerative diseases that have a multifactorial nature and 4-6 discrete molecular steps in their pathogenesis (16, 17). For example, experimental evidence points to a role for vigorous exercise leading to the build-up of reactive oxygen species (18), viral infections (19–21) and stress-induced proteostasis impairment (22), although the exact nature of extrinsic factors that act as “second hits” in ALS is yet to be elucidated. Better understanding of STMN2 regulation under stress may provide novel clues into the pathogenesis of ALS and other neurodegenerative diseases.

Here we report that STMN2 protein is disproportionately affected by stress response, both acute and chronic. Acute stress causes its rapid depletion due to activated proteasome degradation and stress granule-mediated translation repression, preceding cryptic splicing-mediated depletion due to TDP-43 loss-of-function. On the other hand, being an extremely short-lived protein, STMN2 is highly sensitive to mild translation deficits such as those associated with chronic low-grade stress. Analysis of human post-mortem tissue revealed that STMN2 protein level is maintained in non-TDP ALS (FUS and SOD1) – despite reported translation deficits and/or stress granule dysmetabolism in these subtypes. Data from cell models suggested compensatory STMN2 mRNA upregulation as a possible mechanism. Consistent with the ongoing compensation at the RNA level, STMN2 mRNA is also upregulated in the relatively spared CNS areas in ALS-TDP.

Thus, we report novel molecular insights into STMN2 regulation under stress and in neurodegeneration, linking this protein to the common molecular denominators of the latter process – misregulation of translation and stress granule metabolism.

## METHODS

### General cell culture and transfection

Human SH-SY5Y neuroblastoma cells were obtained from ATCC via Sigma and cultured in Dulbecco’s Modified Eagle Medium/Nutrient Mixture F-12 (DMEM/F-12) supplemented with 10% foetal bovine serum (FBS) and penicillin-streptomycin (all Thermo Fisher Scientific). Neuro2a murine cells (gift from Guillaume Hautbergue) were cultured in DMEM/F-12 supplemented with 2 mM Glutamine, 1% Non-Essential Amino Acids (NEAA) and 10% FBS. The cells were seeded onto optical 96-well plates (Phenoplate-96, PerkinElmer) for imaging experiments on Opera Phenix and Sartorius Incucyte S3 or onto 24-well plates (with or without coverslips) for all other experiments. Toxicity analysis was performed using Promega Cell Titer Blue kit according to the manufacturer’s instructions. Plasmids for the expression of GFP-tagged G3BP1 and STMN2 WT/3A (all in pEGFP-N1 vector) were custom-made by Genewiz/Azenta and verified by sequencing. STMN2 siRNA was from ThermoFisher (SilencerSelect®) and scrambled control siRNA was AllStars from Qiagen. Cells were transfected using Lipofectamine2000.

### Motor neuron differentiation

Human motor neurons were differentiated from the KOLF2.1J iPSC line (obtained from iNDI/JAX, JIPSC001000) as described for human ES cells previously (23). Briefly, iPSCs were maintained on Matrigel® (Corning)-coated 6-well plates in mTeSR Plus medium (Stemcell Technologies), with media changes performed daily. When cells reached 90% confluency, they were detached using ReLeaSR® (Stemcell Technologies) and re-plated at the 1:4 ratio using the Rock kinase inhibitor Y-27632 (ApexBio), with a complete media change after 24 hours. When iPSCs reached 70-80% confluency, media was changed to “DM1” differentiation medium composed of Advanced DMEM/F12 (ADF) supplemented with 10 μM SB431542 (Abcam) and 1xB27 (without vitamin A; Gibco). Daily media changes were performed throughout differentiation with supplements added according to the differentiation stage. On day 4, purmorphamine (1 μM, Cayman Chemicals) and retinoic acid (0.1 μM, Sigma) were added to DM1 and cells were cultured in this medium (“DM2”) until day 16. Neural precursors (NPCs) were split at the 1:2 ratio on day 8 and on day 16 using Accutase®. From day 16, NPCs were transferred to “DM3” medium composed of ADF, 1xB27, 1xN2 supplement (Gibco) and 10 ng/ml human BDNF (Miltenyi). On day 23, motor neurons were lifted using Accutase and re-plated in DM3 media. After recovery, the cells were changed into “DM4” media – DM3 with half of ADF replaced with Neurobasal A (Gibco), and matured for at least 7 days, before re-plating in the final (experimental) format.

### Chemical treatments

For acute oxidative stress induction in SH-SY5Y cells, cultures were treated with 500 µM of NaAsO_2_ (Sigma) for 1 hour and washed with fresh media once. Other pharmacological treatments (purchased from Merck/Sigma unless indicated otherwise): MG132 (10 µM), thapsigargin (10 µM), poly(I:C) (100 ng per well), ISRIB (500 nM), lipoamide (10 µM), silvestrol (100 nM), rocaglamide A (500 nM), cycloheximide (10 µg/ml), pitavastatin and simvastatin (both ApexBio; various concentrations). Incubation times within individual experiments and any other modifications to the protocol are indicated in the respective Figure legends and in the Results section. Me4BodipyFL-Ahx3Leu3VS (Proteasome Activity Probe, I-190, R&D Systems) was added to live cells (final concentration of 10 µM) 5 min prior to imaging on Opera Phenix.

### Immunocytochemistry, puro-PLA, and fluorescent microscopy

Immunocytochemistry was performed on cells growing either on 10-mm coverslips (VWR) in 24-well plates or in 96-well optical plates (Phenoplate-96; PerkinElmer). Cells were fixed with 4% PFA in 1xPBS, washed with 1xPBS and kept in 70% ethanol at 4LC. Immunostaining was performed as described earlier (24) using commercially available antibodies (all at 1:1000 dilution): STMN2 (rabbit polyclonal, Proteintech, 10586-1-AP or mouse monoclonal, Proteintech, 67201-1-Ig); TDP-43 (rabbit polyclonal, C-terminal, Sigma or mouse monoclonal, R&D Systems, MAB7778); G3BP1 (rabbit polyclonal, Proteintech, 13057-2-AP); GM130 (rabbit polyclonal, Proteintech 11308-1-AP); Tuj/betaIIItubulin (rabbit recombinant monoclonal Alexa488-labelled, Abcam, AB237350). For puromycin labeling coupled with proximity ligation assay (puro-PLA), cells were cultured in optical plates. Puromycin (Sigma, 1 µg/ml) was added directly to culturing media for the indicated time, and cells were fixed as above. Fixed cells were washed with 1xPBS and incubated in a combination of primary antibodies (1:1000) against puromycin (mouse monoclonal, Merck, MABE343) and STMN2 (rabbit polyclonal, Proteintech) diluted in PBST, overnight at 4LC. PLA was performed using Duolink® In Situ Orange Starter Kit Mouse/Rabbit (Merck/Sigma). Nuclei were stained with DAPI. Conventional fluorescence microscopy of immunostained cells was performed using a 100x oil objective on an Olympus BX57 upright microscope equipped with ORCA-Flash 4.0 camera (Hamamatsu) and cellSens Dimension software (Olympus). High-content microscopy in the 96-well format was performed on Opera Phenix and Harmony 4.9 software.

### RNA in situ hybridisation

RNA-FISH was performed using a custom STMN2 mRNA probe set (24 probes, Supplementary Table S1) designed and synthesised by Biosearch Technologies, according to the previously described protocol (24). BaseScope ISH was performed with the probes for full-length STMN2 mRNA – 1048241-C1 (exon 1) or 1048231-C1 (exons 4 and 5) – according to the manufacturer’s instructions. Briefly, cells grown on coverslips were fixed with 10% neutral buffered formalin for 30 min at RT on a flat-surface shaker. Cells were then gradually dehydrated in ethanol with ascending concentration for 5 min each and kept in 100% ethanol at -20°C. Coverslips were rehydrated by ethanol washes with descending concentration for 5 min each, before 1xPBS was added to coverslips for 10 min. Samples were then treated with 30 µl protease III (ACDBio) at RT for 15 min. Coverslips were washed again with fresh 1xPBS, before incubating in 30 µl of a BaseScope probe for 2 hours at 40°C in HybEZ oven. Signal was detected using BaseScope Detection kit, according to manufacturer instructions, after which Fast Red substrate was added for 10 min at RT. Coverslips were mounted onto glass slides with Immu-Mount (ThermoFisher).

### Image analysis and quantification

Manual quantification of RNA-FISH experiments was performed using custom pipelines on Image J. Automated quantification of stress granules was carried out using custom pipelines (Spot analysis) on Harmony 4.9 software as described previously (25). Automated quantification of puro-PLA signals was performed using a custom Spot analysis pipeline on Harmony 4.9. For the analysis with the proteasome activity probe, fluorescence intensity (GFP channel) was measured in individual cells using Harmony 4.9 software.

### Longitudinal analysis of cell viability on Incucyte

SH-SY5Y cells or Day 32 human motor neurons were plated onto optical 96-well plates (Phenoplate-96). Motor neurons were matured directly on plates for 7 days before the analysis. Viability and proliferation analysis were performed on Incucyte® S3 Live-Cell Analysis System (Sartorius), by recording cell density in the bright-field (Phase Object Count) and apoptosis rate using the Incucyte® Caspase-3/7 Green Dye (Green Object Count).

### RNA expression analysis

Total RNA was extracted using GenElute total mammalian RNA kit (Sigma) or QiaZOL (Qiagen) in accordance with the manufacturer’s instructions. First-strand cDNA synthesis was performed using 500 ng of RNA with random primers (ThermoFisher) and MMLV reverse transcriptase (Promega). qRT-PCR was performed using qPCRBIO SyGreen Lo-ROX (PCRbio), and GAPDH was used for normalisation. Human-specific primers from our previous study were used (15), and STMN2 pre-mRNA primers were as follows: 5’-CTCGGCAGAAGACCTTCGAG-3’ and 5’-ACAAGCCGCATTCACATTCA-3’. Mouse-specific primers were as follows: Stmn2, 5’-CTCCTCATGGATTACGCGCT-3’ and 5’-TCAGCACGTTGGAAGGTTCA-3’; Gapdh, 5’-AGGTCGGTGTGAACGGATTTG-3’ and R 5’-GGGGTCGTTGATGGCAACA-3’.

### In vitro protein assays

For the analysis of STMN2 phosphorylation, cells were lysed in the lysis buffer (50mM Tris HCl, 150mM NaCl, 1% Triton X100, protease inhibitor cocktail), scraped, and suspension was vortexed every 5 min for 30 min for complete cell lysis. Lysate was mixed with 2x Calf Intestinal Alkaline Phosphatase (CIP) in reaction buffer (50mM NaCl, 50mM Tris-HCl, 20mM MgCl_2_, 2mM DTT, protease inhibitor cocktail), based on 1 unit per 1 µg protein (or the same volume of water as control). Samples were incubated with CIP at 37°C for 1 hour and analysed by western blot. For the analysis of STMN2 solubility under stress, samples were incubated in the urea lysis buffer – 50mM Tris, 150mM NaCl, 0.5% Triton-X, 8M urea, before mixing with 2xLaemmli buffer and proceeding to western blot. For puromycilation (measurement of protein translation rates), cells were pulsed with 1 µM puromycin for 10 min prior to lysis.

### Western blotting

Cells were lysed in 2xLaemmli buffer, and the samples were heated at 95°C for 15 min. Samples were resolved on a 10% Mini-PROTEAN® TGX™ hand-cast protein gel (Bio-Rad) and transferred to a PVDF membrane (Amersham). Gels were stained with Gelcode^TM^ (ThermoFisher) post-transfer for total protein (loading control). Membranes were blocked for 1 hour in 4% non-fat milk/TBST and then incubated with primary antibodies (1:500 for puromycin and 1:1000 all others) at 4℃ overnight: STMN2 (rabbit polyclonal, Proteintech, 10586-1-AP); TDP-43 (rabbit polyclonal, C-terminal, Sigma); p-eIF2α (rabbit monoclonal, Abcam, ab32157); total eIF2α (rabbit polyclonal, Cell Signaling, 9722), and puromycin (mouse monoclonal, Merck, MABE343). Anti-rabbit or anti-mouse HRP secondary antibody (GE Healthcare/Amersham) incubation was performed for 1 hour at RT in 4 % milk/TBST. Signal was detected using Clarity Max Western ECL Substrate (Bio-Rad) and quantified using Licor Odyssey FC/Image Studio software or Image J.

### Analysis in human tissue

Spinal cord sections from neuropathologically characterised individuals from the Sheffield Brain Tissue Bank (SBTB) were used (see Table 1). SBTB has ethical permission to function as a research tissue bank from the Wales Research Ethics Committee 5 (Reference 24/WA/0052). SBTB adheres to consenting protocols laid down by the UK Human Tissue Authority and agreed to by the Research Ethics Committee. Sections (7 µm thick) were processed for immunohistochemistry as described before (26), except antigen retrieval in 10 mM sodium citrate (pH 6.0) was performed in a pressure cooker and TBS buffer was used for the washes. Sections were immunostained for STMN2 using a rabbit polyclonal antibody (Proteintech, 10586-1-AP) or mouse monoclonal antibody (Proteintech, 67201-1-Ig), with ImPACT® DAB reagent kit used for signal detection. Slides were covered with coverslips using DPX. BaseScope ISH analysis on rehydrated sections was performed as described above for cells on coverslips; 1048231-C1 probe was used. Imaging was performed either on Hamamatsu NanoZoomer XR slide scanner or on Nikon Eclipse microscope. Image analysis (pixel intensity in individual neurons) was performed using QuPath v0.5.1 software. qRT-PCR analysis in the motor cortex samples was carried out as described above for cells.

**Table 1.** Human cases used in the study.

| ID | Type | Sex | Age at death |
| --- | --- | --- | --- |
| **Immunohistochemistry** |  |  |  |
| NA014/1999 | Control (group 1) | F | 86 |
| NA019/1991 | Control (group 1) | M | 54 |
| NA025/1995 | Control (group 1) | M | 67 |
| LP085/2007 | Control (group 1) | F | 59 |
| LP005/2007 | Control (group 1) | M | 63 |
| LP005/2011 | sALS | F | 76 |
| LP009/2008 | sALS | M | 76 |
| LP014/2011 | sALS | M | 52 |
| LP024/2008 | sALS | F | 63 |
| LP026/2013 | sALS | M | 62 |
| 098/2007V | Control (group 2) | M | 68 |
| 135/2018 | Control (group 2) | F | 85 |
| 005/2007 | Control (group 2) | M | 63 |
| PA000157V/23 | FUS | M | 21 |
| 1044/1998 | FUS | M | 62 |
| LP024/2009 | FUS | M | 72 |
| 037/2004 | SOD1 | F | 66 |
| 034/1992 | SOD1 | F | 37 |
| **RNA expression analysis** |  |  |  |
| 071/1992 | Control (group 3) | F | 76 |
| 025/1995 | Control (group 3) | M | 67 |
| 178/1995 | Control (group 3) | F | 63 |
| 293/1991 | Control (group 3) | F | 65 |
| 042/2005 | sALS | M | 65 |
| 039/2005 | sALS | F | 61 |
| 075/2008 | sALS | F | 61 |
| 049/2005 | sALS | F | 64 |
| 063/08 | ALS-C9 | F | 69 |
| 046/10 | ALS-C9 | F | 67 |
| 041/04 | ALS-C9 | M | 64 |
| 098/02 | ALS-C9 | F | 64 |

### NYGC ALS consortium RNA-seq dataset analysis

NYGC cohort RNA sequencing data was obtained from GEO omnibus (GEO accession no. GSE137810). Raw counts were converted to approximate RPKM values using gene length estimations and per-sample library size scaling. Counts corresponding to STMN2 mRNA were identified by filtering for Ensembl gene identifier ENSG00000104435. ALS samples were then classified as TDP-related ALS, ALS-SOD1 and ALS-FUS based on the “Subject Group” metadata field, and TPM for STMN2 plotted individually for motor cortex, frontal cortex and spinal cord tissues. Statistical comparisons between ALS and control samples were performed using limma (27) as previously described (28).

### Statistical analysis

Datasets were tested for normality using appropriate tests in GraphPad Prism (version 10.6.1). P-values were calculated in GraphPad Prism using an appropriate parametric or non-parametric test, depending on the nature of the data and the recommended post-hoc test for correction for multiple comparisons, where applicable. Experiments were repeated at least three times, with similar results. N indicates the number of biological replicates (obtained in independent experiments). Error bars represent S.E.M. unless indicated otherwise.

## RESULTS

### STMN2 protein depletion late in stress is caused by TDP-43 loss of function

We previously found, using recovery from a reversible stressor sodium arsenite (NaAsO_2_) as a cell stress paradigm, that STMN2 depletion correlates with nuclear condensation of TDP-43, loss of its solubility and activation of cryptic splicing (15). Here, we first confirmed stress-induced STMN2 depletion in human motor neurons – previously demonstrated by qPCR and western blot (15) – using BaseScope® RNA *in situ* hybridisation with STMN2 full-length mRNA probes and immunocytochemistry (Fig. 1a,b; Fig.S1a). In the experiments with NaAsO_2_-induced stress and recovery, cells were treated with 500 µM NaAsO_2_ for 1 hour and left to recover in the absence of stressor, unless indicated otherwise. We next investigated whether other stresses associated with nuclear TDP-43 condensation (15) lead to STMN2 protein depletion. The ER stressor thapsigargin and viral infection mimic polyinosinic:polycytidylic acid (p(I:C)) led to nearly complete STMN2 protein loss 3 hours post-treatment or 8 hours post-transfection, respectively (Fig.1c). A delayed response to p(I:C) was likely due to its delivery by lipofection, requiring more time to build a stress response. In contrast, treatment with a proteasome inhibitor MG132 led to STMN2 protein accumulation (Fig.1c), consistent with its proteasome-mediated turnover (29). To test the efficiency of STMN2 clearance under stress, we employed treatment with statins that upregulate stathmins via inhibition of the mevalonate pathway (Nolan et al., submitted; Fig.S1b). SH-SY5Y cells were treated with low and high concentrations of statins simvastatin or pitavastatin for 16 hours prior to arsenite stress. Despite substantial STMN2 upregulation in statin pre-treated cells, especially with higher statin concentrations, the protein was still almost completely cleared after 6 hours of recovery (Fig.1d; Fig.S1c).

**Fig. 1.**
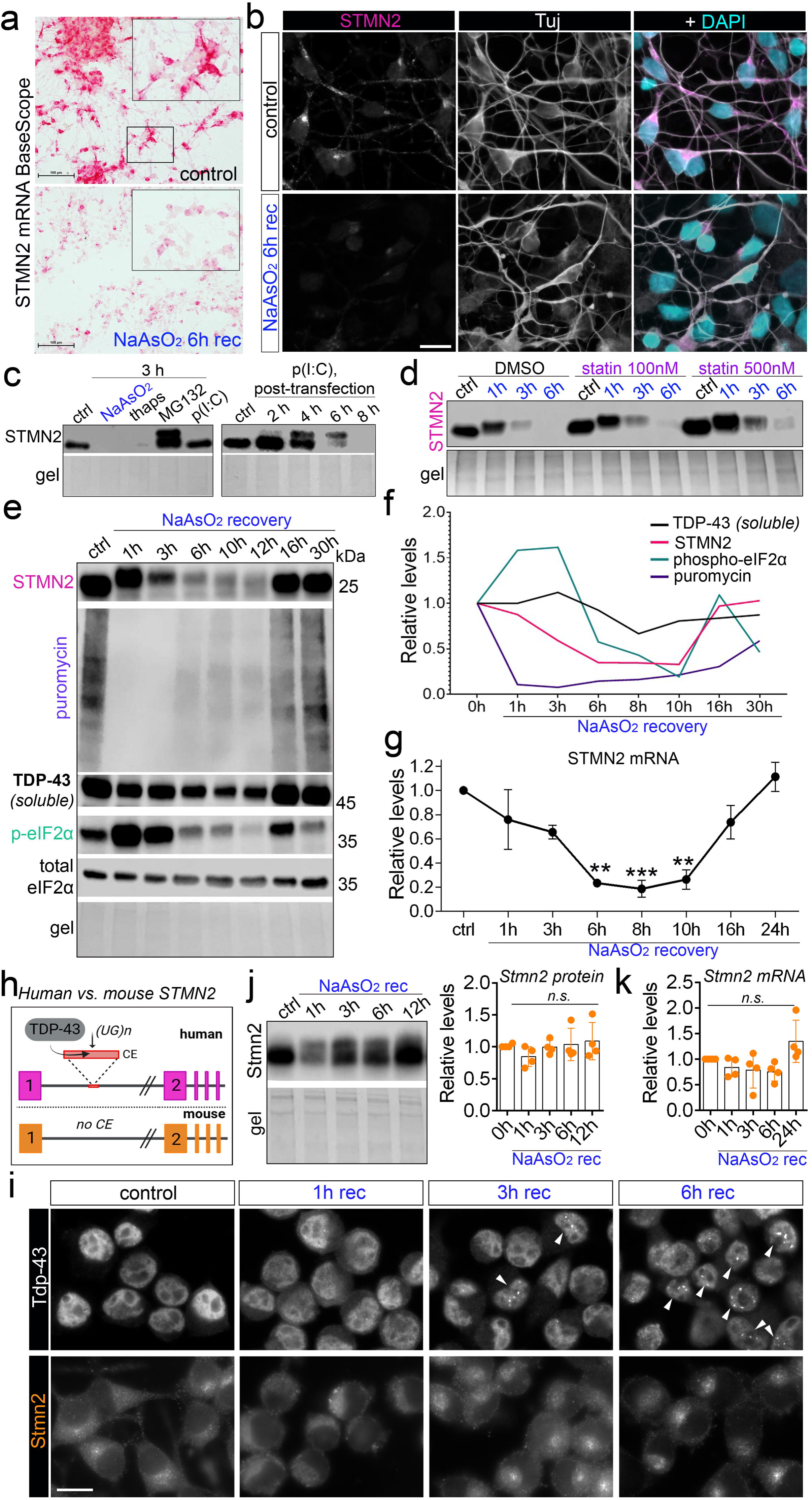
STMN2 protein depletion late in stress is due to TDP-43 loss of function. **a,b** STMN2 mRNA depletion in human motor neurons late in stress/recovery visualised with BaseScope ISH (a) and immunostaining (b). Probe #1048241-C1 was used in a. Representative images are shown. Scale bars, 100 μm in a and 10 μm in b. **c** Cellular stressors with different mechanism of action cause STMN2 protein depletion. Representative western blots are shown. **d** Statin pre-treatment upregulates STMN2 protein but does not prevent its stress-induced clearance. Cells were pre-treated with pitavastatin for 16 h. Time of recovery from NaAsO_2_ stress is indicated. **e,f** Analysis of STMN2 protein dynamics and translational repression during the recovery from NaAsO_2_ stress. Nascent protein production was analysed by ribopuromycilation. Representative western blot (e) and its quantification (f) are shown. **g** STMN2 mRNA depletion and subsequent restoration correlate with STMN2 protein level changes during stress recovery. N=3, **p<0.01, ***p<0.001, one-way ANOVA with Dunnett’s post-hoc test. **h** STMN2 splicing regulation in human and mouse cells. Schematic was prepared using BioRender. **i** Tdp-43 protein undergoes recruitment into nuclear condensates, however Stmn2 protein levels and nuclear-cytoplasmic distribution remain unaffected during the recovery from NaAsO_2_ stress in Neuro2A cells. Representative images are shown. Arrowheads indicate cells with nuclear TDP-43 condensates. Scale bar, 10 μm. **j,k** Stmn2 protein (j) or mRNA (k) levels do not significantly change under cellular stress in mouse Neuro2A cells. Representative western blot and quantification are shown in j. mRNA levels in k were measured using qRT-PCR. N=4, n.s., non-significant. SH-SY5Y cells were used in c-g.

We next performed a detailed analysis of the temporal dynamics of STMN2 mRNA and protein levels and TDP-43 solubility across 30 hours of recovery from NaAsO_2_ stress, in SH-SY5Y cells. STMN2 protein was almost completely depleted after 6 hours of recovery, yet its level was fully restored between 12 and 16 hours of recovery (Fig. 1e,f). STMN2 protein level correlated well with STMN2 mRNA level (Fig. 1g) and with TDP-43 solubility (Fig. 1e), where STMN2 mRNA was reduced to <20% of the steady-state level between 6 and 10 hours of recovery (Fig. 1g). However, ribopuromycilation assay revealed a global shutdown of protein synthesis (loss of puromycin signal corresponding to its incorporation into nascent peptides) between 1 and 3 hours of recovery, with its gradual restoration from 6 h of recovery onward (Fig. 1e,f). Consistently, phospho-eIF2α, a marker of inhibited translation, was dramatically upregulated in this time window (Fig. 1e,f). Therefore, STMN2 protein loss could be due to stress-induced translational repression.

To address the primary mechanism directly, we used a murine neuroblastoma cell line, Neuro2a (N2a) – since mouse *Stmn2* gene lacks the TDP-43-regulated cryptic exon (CE) (Klim et al., 2019) (Fig. 1h). First, we confirmed that mouse Tdp-43 also forms nuclear condensates during the recovery from NaAsO_2_ stress, with temporal dynamics similar to human cells (detectable from 3 hours and peaking at 6 hours of recovery; Fig. 1i). We next demonstrated that both Stmn2 protein and mRNA levels were sustained during stress and recovery in mouse cells (Fig. 1j,k). Therefore, human-specific STMN2 depletion late in stress is primarily due to TDP-43 loss of function and activated cryptic splicing rather than attenuated global translation. Indeed, despite partial restoration of global translation (as indicated by puromycin incorporation) from 6 hours of recovery, STMN2 protein levels continued to decline (Fig. 1e,f), which coincided with STMN2 mRNA depletion (Fig. 1g).

### STMN2 protein depletion early in stress is caused by proteasomal degradation and phosphorylation

To establish the primary route of STMN2 protein clearance during stress, SH-SY5Y cells were treated with the proteasome inhibitor MG132 or autophagosome-lysosome inhibitor chloroquine (CQ), prior to NaAsO_2_ addition. MG132 but not CQ prevented STMN2 depletion between 1 and 6 hours of NaAsO_2_ stress recovery (Fig. 2a). Using a proteasome activity probe Me4BodipyFL-Ahx3leu3VS that detects proteolytically active proteasome subunits (30), we found that proteasome activity increases early during stress (30 min of NaAsO_2_), before declining during the recovery period (45 min into the recovery) (Fig. 2b). Consistent with this result, we observed significant STMN2 protein depletion as early as 20 min into stress exposure (Fig. 2c,d). Interestingly, STMN2 protein lost its clustered localisation and became diffusely redistributed in the cytosol from 20 min of stress onward (Fig. 2d). Upon translation block with cycloheximide, to uncouple protein synthesis and degradation, STMN2 protein level at 1 hour of recovery was higher in stressed compared to non-stressed cells (Fig. S2a), supporting that proteasome activity declines at this time-point. MG132 pre-treatment also prevented STMN2 depletion at these early time-points (Fig. S2b).

**Fig. 2.**
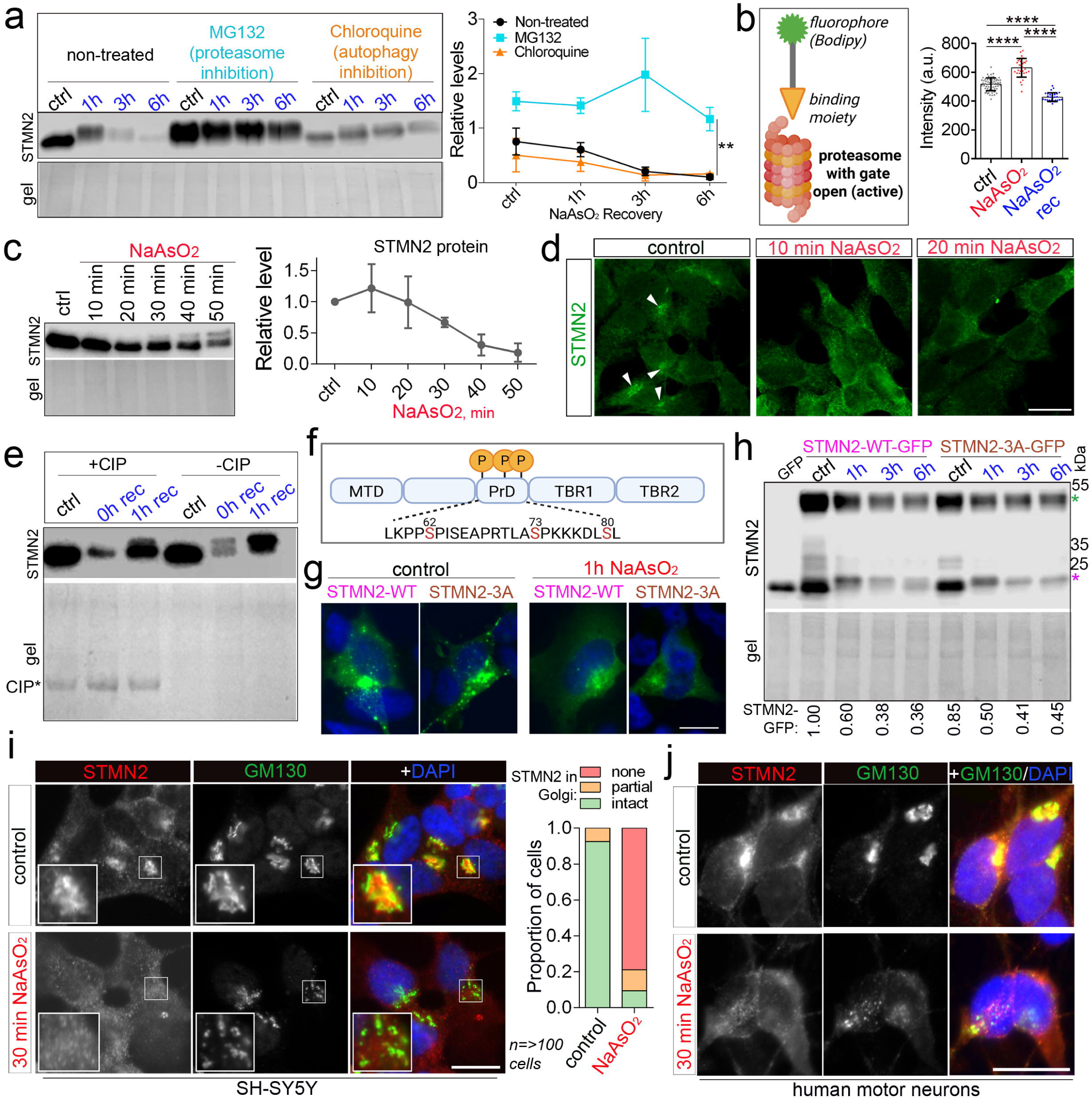
Mechanisms of STMN2 regulation early during stress. **a** STMN2 protein depletion during stress is mediated by the proteasome. Cells were pre-treated with the compound for 1 hour prior to arsenite stress and were cultured in the presence of the compound during the recovery from arsenite. Representative western blot and quantification are shown. N=3, **p<0.01, one-tailed paired *t* test. **b** Activation of proteasomal degradation early in stress and its subsequent decline during the recovery from stress. Proteasome activity probe was added directly to the media before live imaging on Opera Phenix. Fluorescence intensity was plotted from individual cells. N=3, n=30; ****p<0.0001, one-way ANOVA with Tukey post-hoc test. **c,d** STMN2 protein becomes redistributed and depleted within minutes of NaAsO_2_ addition. Representative western blot and its quantification (c) and representative immunostained images (d) are shown. N=3. Arrowheads indicate membrane-associated STMN2 protein. **e** STMN2 undergoes phosphorylation early during stress. Calf intestinal phosphatase (CIP) treatment of cell lysates reverses stress-induced STMN2 mobility shift. Representative western blot is shown (experiment repeated 3 times). **f** STMN2 phosphorylation sites. STMN2-3A: three Serines (in red) were mutated to Alanines. MTD, membrane-targeting domain; PrD, proline-rich domain; TBR, tubulin-binding repeat domain. **g,h** Reduced phosphorylation of STMN2 does not affect its distribution or stability. Representative immunostaining images (g) and western blot (h) are shown. Purple asterisk indicates endogenous STMN2 and green asterisk – GFP-tagged variants. Protein levels were quantified by densitometry. N=3. **i,j** STMN2 redistribution correlates with Golgi fragmentation early in stress. Representative images and quantification for SH-SY5Y cells (i) and representative images for human motor neurons (j) are shown. “None”, “partial”, and “intact” – STMN2 overlap with Golgi staining. SH-SY5Y cells were used except in j. Scale bars, 10 μm. Schematics in b and f were prepared using BioRender.

In addition to its gradual depletion, STMN2 protein underwent a shift to a slower migrating band from 50-60 min of NaAsO_2_ stress, suggestive of a post-translational modification (PTM) (Fig. 2a,c). This shift was also detectable with other stressors (Fig. 1c). Stathmins are phosphoproteins and can be phosphorylated under various physiological conditions, which leads to reduced binding to tubulin and loss of function in tubulin sequestration (31). Phosphorylation has been associated with a functionally inactive STMN2 state (32). Treatment of SH-SY5Y cell lysates from control and stressed cells with calf intestinal alkaline phosphatase (CIP) reversed stress-induced STMN2 mobility change (Fig. 2e), confirming that STMN2 undergoes extensive phosphorylation under stress. We next asked whether this PTM influences stress-induced STMN2 clearance. We generated constructs to express GFP-tagged WT STMN2 and its non-phosphorylatable form (3A) – with three serines in the proline-rich domain (PrD) mutated to alanines, including the two most frequently phosphorylated ones (Ser-62, Ser-73) (33) (Fig. 2f). STMN2-3A variant was expressed and distributed similar to WT protein, with the typical clustered localisation (Fig. 2g), and loss of phosphorylation on these sites did not affect the rate of STMN2 clearance during stress (Fig. 2h).

STMN2 prominently localises to the Golgi apparatus membranes (34). Using a Golgi marker GM130, we confirmed that STMN2 loses its localisation to Golgi early during stress (30 min of arsenite treatment), including in human motor neurons, which coincides with Golgi fragmentation (Fig. 2i,j).

Thus, STMN2 is subject to extremely rapid and drastic changes in its abundance, localisation and functional state early during stress (on the scale of minutes), which precedes the activation of TDP-43-regulated cryptic splicing.

### STMN2 translation remains active under acute stress

We made an unexpected observation that despite the dramatic depletion of STMN2 protein after 30-60 min of arsenite stress in SH-SY5Y cells or human motor neurons, it is restored to near-normal level after 1 hour of recovery (2 hours post-arsenite addition in total), before being depleted again (Fig. 3a,b; Fig.S2c). This effect was not due to fluctuations in STMN2 solubility, since its transient depletion was still detectable after urea extraction (Fig.S2d). This phenotype was confirmed using an independent (mouse monoclonal) STMN2 antibody (Fig. S2e). These experiments suggested that STMN2 is actively translated during stress despite global translational shutdown. STMN2 mRNA is relatively stable, and its level is maintained early in stress (Fig. 1g), which should enable translation. To selectively measure STMN2 translation, we employed puro-PLA – proximity ligation assay coupled with puromycin labelling, which allows quantitative detection of individual protein molecule synthesis *in situ* (35) (Fig. 3c). STMN2 translation could be readily detected under basal conditions (>100 PLA signals per cell with a 10-min puromycin pulse), but was completely abolished by cycloheximide co-treatment and was severely reduced after 1 hour of stress (Fig. 3d; Fig. S2f). However, accumulation of the puro-PLA signal for 30 min during the recovery (continuous puromycin presence in the media between 10 and 40 min of recovery), allowed the detection of ongoing STMN2 translation – which remained at ∼30% of the basal level (Fig. 3d). S*TMN2* gene sequence analysis (for putative IRES and other elements) failed to identify any regulatory elements that may underlie active STMN2 translation during stress (data not shown). Interestingly, STMN2 mRNA lacks a Kozak consensus sequence. Efficient translation of STMN2 mRNA under stress may be possible due to its high abundance (in the top 25 mRNAs in neurons). Notably, when the proteasome was inhibited in stressed cells, STMN2 accumulated at a level higher than in non-stressed cells (Fig. S2b; compare the 1-hour or 3-hour recovery time-points with and without MG132), indicating its ongoing translation under the conditions of declining proteasome activity.

**Fig. 3.**
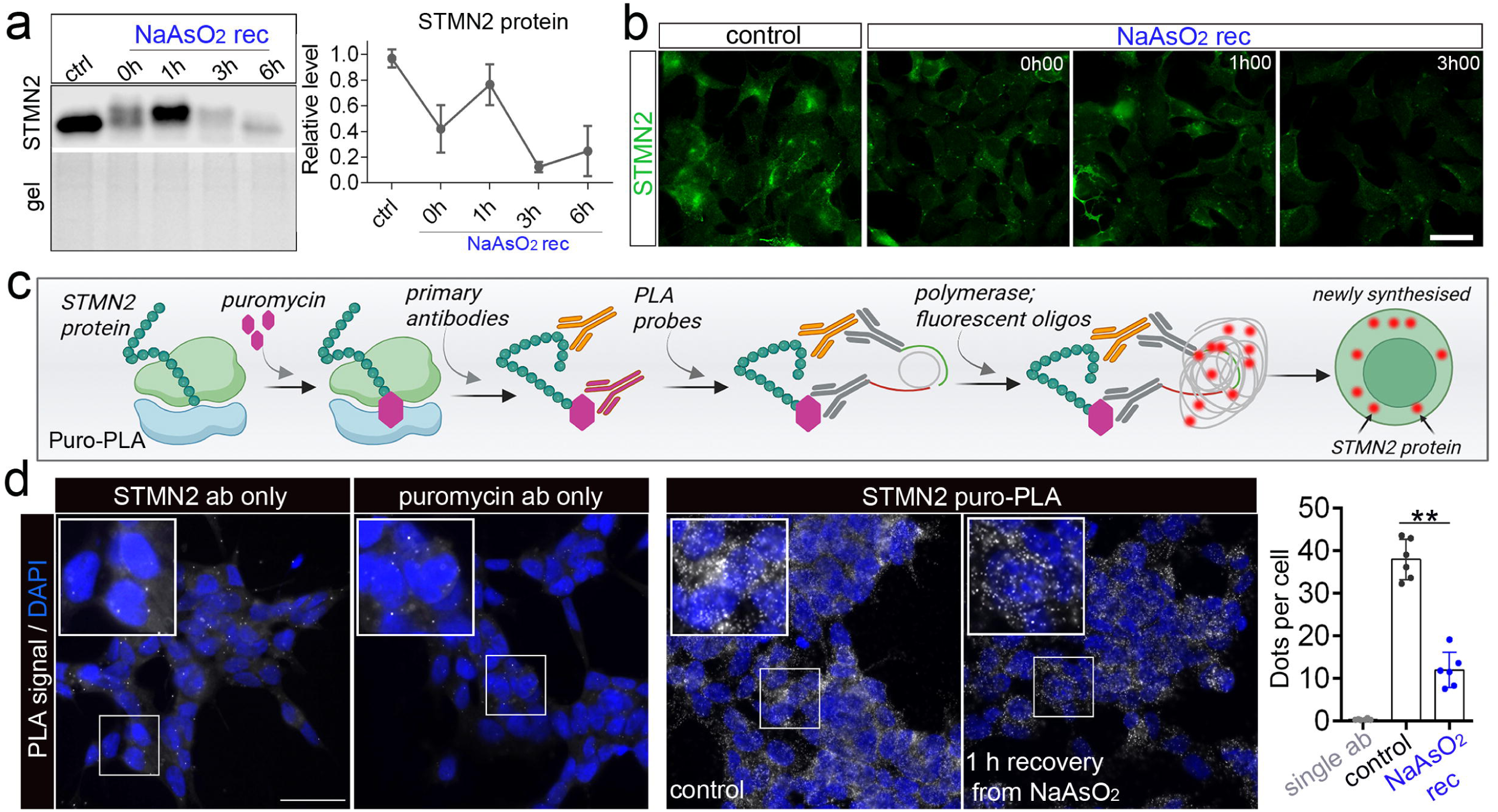
STMN2 translation can be maintained under acute stress characterised by global translational repression. **a,b** Transient restoration of STMN2 protein level early during the recovery from arsenite stress. Representative western blot and quantification (a) and representative images for immunostaining (b) are shown. N=3-6. Error bars represent S.D. Scale bar, 10 μm. **c** Puro-PLA approach. Schematic was prepared using BioRender. **d** Ongoing STMN2 translation under acute stress, as revealed by puro-PLA analysis. Puro-PLA signals were quantified in an automated way on Opera Phenix/Harmony. N=6, n>3000 cells per condition. **p<0.01, Mann-Whitney *U* test. “One ab”, single antibody was used (control condition). Scale bar, 25 μm. SH-SY5Y cells were used.

### STMN2 translation is limited by stress granules

We noticed that STMN2 undergoes rapid depletion in chloroquine (CQ)-treated cells in the absence of NaAsO_2_ stress – being dramatically downregulated after just 1 hour of pre-treatment (Fig. 2a). Similar to other chemical treatments examined here (NaAsO_2_, thapsigargin, p(I:C)), CQ induces eIF2α phosphorylation typical for the activation of integrated stress response (36). This led us to hypothesise that STMN2 downregulation is triggered by treatments associated with translational repression. To test this, we examined whether stressors that repress translation via an alternative mechanism would also affect STMN2 protein (Fig. 4a). Small molecule inhibitors of another translation initiation factor, eIF4A, silvestrol and rocaglamide-A (37) both caused dramatic depletion of STMN2 within only 1 hour of treatment, being more efficient than phosho-eIF2α-dependent stressors sorbitol and thapsgargin included as positive controls (Fig. 4b,c).

**Fig. 4.**
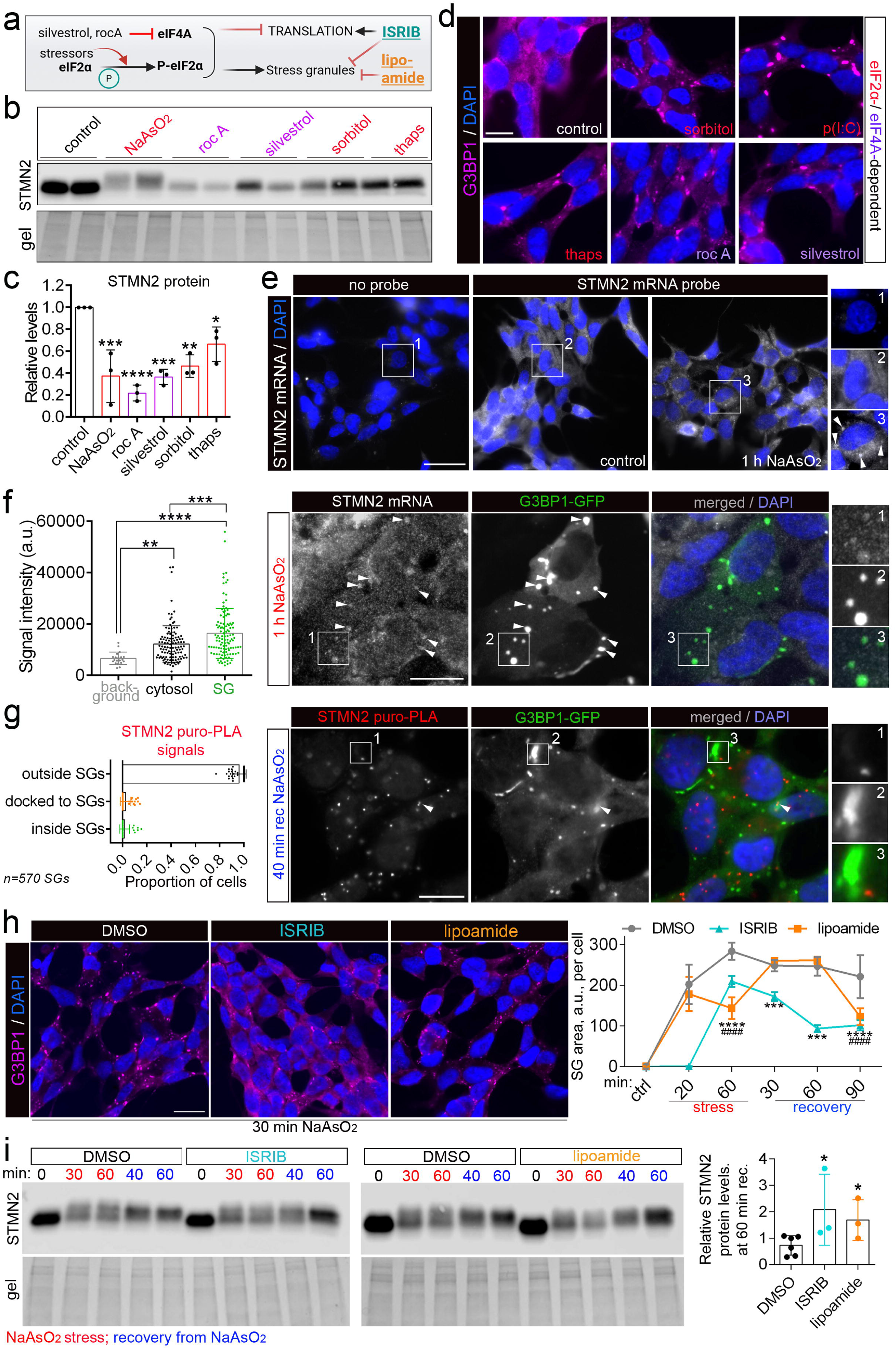
Stress granules contribute to STMN2 regulation under stress. **a** Stress granule assembly and translational repression under stress and their modulation by small molecules. Schematic was prepared using BioRender. **b,c** eIF4A inhibition with small molecules leads to rapid and dramatic STMN2 depletion. Representative western blot (b) and quantification (c) are shown. Cells were treated with the indicated compounds for 1 hour. N=3; *p<0.05, **p<0.01, ***p<0.001, ****p<0.0001, one-way ANOVA with Dunnett’s post-hoc test. **d** eIF4A inhibitors and phospho-eF2α dependent stressors induce stress granule assembly. Representative images are shown. Scale bar, 10 μm. **e** STMN2 mRNA segregates into distinct foci in response to stress. Single molecule RNA-FISH with a custom probe was used. Arrowheads point to the foci with STMN2 mRNA enrichment. Scale bar, 25 μm. **f** STMN2 mRNA is significantly enriched in stress granules. Quantification and representative images are shown. Arrowheads point to G3BP1-positive foci – stress granules (SGs). Stress granules from 114 cells were included in the analysis. **p<0.01, ***p<0.001, ****p<0.0001, one-way ANOVA with Dunnett’s post-hoc test. Scale bar, 10 μm. **g** STMN2 translation occurs outside stress granules. Newly synthesised STMN2 protein molecules were visualised by puro-PLA and SGs – by G3BP1 immunocytochemistry. 570 stress granules were included in the analysis. Scale bar 10 µm. **h** ISRIB and lipoamide interfere with stress granule assembly. NaAsO_2_ was used. Representative images and automated quantification data are shown. N=3 (500-1000 cells analysed in each experiment). ***/^###^p<0.001, ****/^####^p<0.0001, two-way ANOVA. Scale bar 10 µm. **i** ISRIB and lipoamide attenuate STMN2 depletion during the recovery from stress. Cells were pre-treated with either ISRIB or lipoamide, which was replenished before the recovery from arsenite (continuous presence of the compound in the media). Representative western blots and quantification are shown. N=3; *p<0.05, Mann-Whitney *U* tests. SH-SY5Y cells were used.

All of the above treatments are known to be associated with the assembly of stress granules (SGs) (38) – RNP condensates thought to be involved in translational repression and dysregulated in neurodegeneration (39) (Fig. 4a,d). We hypothesised that SGs contribute to the regulation of STMN2 protein levels under stress. First, we employed single-molecule (sm)RNA-FISH to study STMN2 mRNA enrichment in SGs. In line with high STMN2 abundance in neuronal cells, high-density mRNA signal was detected, with individual mRNA molecules barely distinguishable; visible signal segregation into foci was observed in NaAsO_2_-treated cells (Fig. 4e). For quantification of STMN2 mRNA enrichment in SGs, these structures were visualised using ectopic expression of G3BP1-GFP. Quantification of STMN2 mRNA signal intensity within SGs and in SG-free cytosol demonstrated that STMN2 mRNA is, on average, 1.32-fold enriched in SGs compared to the surrounding cytosol (Fig. 4f). Although SGs are generally believed to repress translation of the sequestered mRNAs (40), some studies indicate that translation in SGs is not uncommon (41). We used puro-PLA in G3BP1-GFP expressing cells at the 1 hour recovery time-point (when STMN2 protein level is high) to track the origin of the newly synthesised STMN2 protein molecules. STMN2 puro-PLA signals were found almost exclusively outside SGs (Fig. 4g), indicating that STMN2 mRNA translation is inhibited within SGs, consistent with the canonical role for SGs in translational repression. Notably however, a substantial number of STMN2 mRNA molecules were found outside microscopically visible SGs (cytosol vs. background, Fig. 4f). Presumably, this non-SG pool is responsible for STMN2 translation under stress (Fig. 3d).

To directly address the role of SGs in limiting STMN2 translation under stress, we utilised two small molecule inhibitors of SG assembly, ISRIB (42) and lipoamide (43). ISRIB acts downstream of phospho-eIF2α-mediated stress signaling, by increasing active eIF2B subunit assembly and reversing the effects of the eIF2α phosphorylation event; SG assembly is inhibited as a result (44). In contrast, lipoamide directly attenuates SG condensation by changing the redox state of RNA-binding proteins – SG components. Longitudinal analysis with automated quantification demonstrated that treatment with either compound reduces NaAsO_2_-induced SG assembly in SH-SY5Y cells, with ISRIB being more potent than lipoamide (Fig. 4h; Fig. S3a-c). Western blot and immunostaining revealed that ISRIB and lipoamide significantly promote STMN2 protein accumulation during the recovery from NaAsO_2_ stress (Fig. 4i; Fig S3d).

### STMN2 expression is sensitive to chronic translation deficits

The above data suggested that STMN2 translation can be sustained during acute stress, reflecting the vital importance of this protein for neurons. However, the effect of chronically impaired protein translation typical for neurodegenerative diseases (45, 46) remained unclear. To model persistent yet mild translation deficits, we employed ultra-low concentrations of arsenite. Alongside this, we included a repetitive stress paradigm, where acute stress pulses were followed by periods of recovery (Fig. 5a). For repetitive stress, reduced arsenite concentration and duration of treatment, as compared to the standard acute stress, were used, to ensure adequate cell survival for 48 hours (Fig. 5a). STMN2 protein depletion was confirmed under these conditions (Fig. S4a). Although STMN2 protein level was fully restored between stress pulses in the repetitive stress paradigm, the protein was significantly depleted in the chronic stress paradigm both in SH-SY5Y cells and human motor neurons – detectable already at the 24-hour time-point (Fig.5b,c; Fig.S4b,c). Chronic stress did not cause visible nuclear TDP-43 condensation, loss of solubility or cytoplasmic redistribution (Fig. 5b,d). Nor did it cause SG assembly or significantly affect STMN2 mRNA levels (Fig.S4d and not shown). Instead, chronic stress led to attenuated translation and elevated phospho-eIF2α (Fig. 5e; Fig.S4b). CHX pulse-chase demonstrated extremely short STMN2 protein half-time in SH-SY5Y cells – ∼50 min (Fig. 5f). Although more stable in neurons, STMN2 was still short-lived, with a half-life of ∼90 min (Fig. 5f). Therefore, STMN2 protein depletion under chronic stress is likely caused by protein translation deficits that severely impact this extremely short-lived protein, independent of TDP-43 pathology and SGs.

**Fig. 5.**
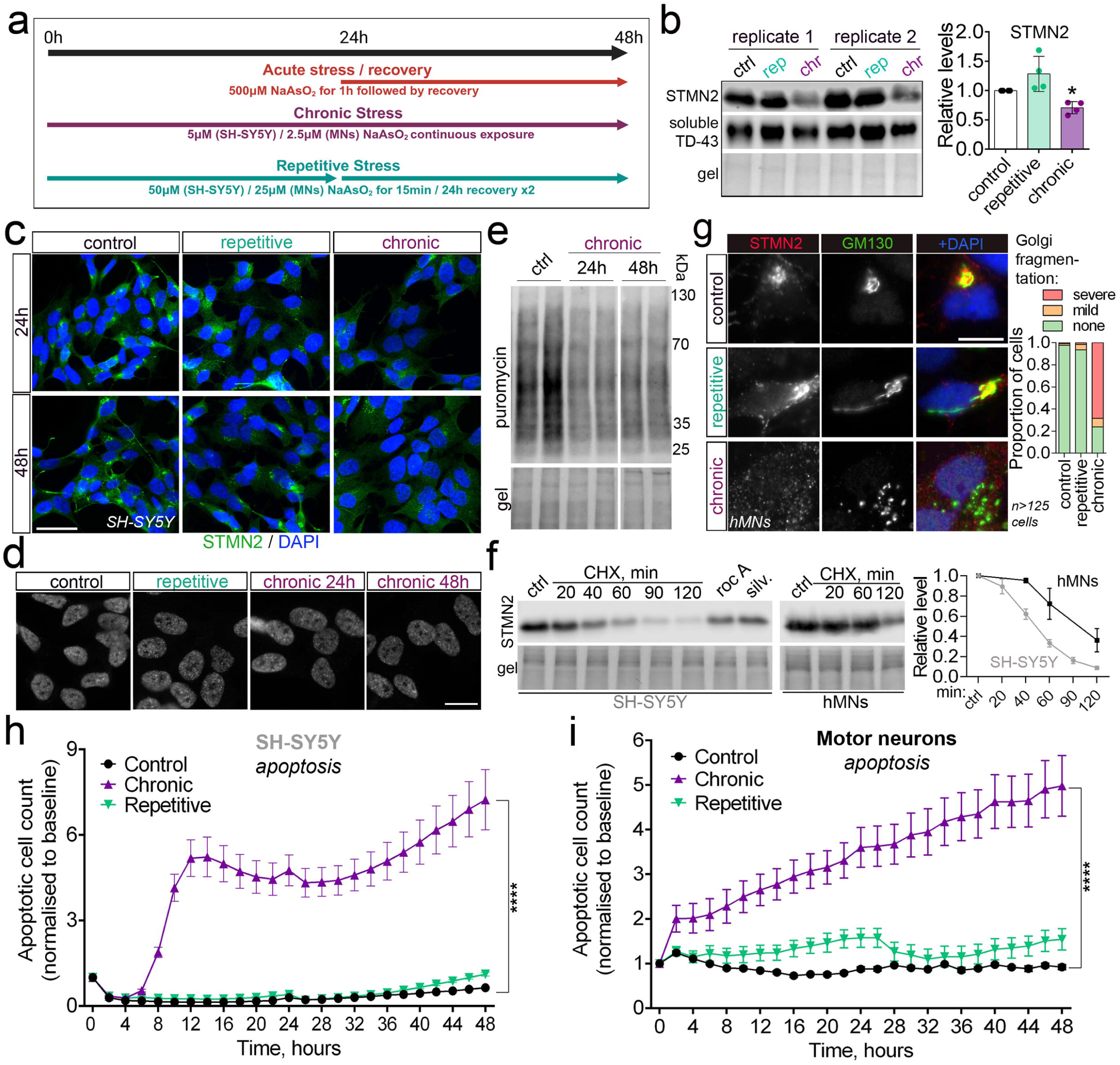
Chronic stress leads to translation deficits and STMN2 protein depletion. **a** Stress paradigms used in the study for SH-SY5Y cells and human motor neurons. Schematic was prepared using BioRender. **b,c** Chronic but not repetitive oxidative stress downregulates STMN2 protein. Representative western blot and quantification (b) and representative immunostaining images (c) are shown for SH-SY5Y cells. N=3; *p<0.05, Mann-Whitney *U* test. Scale bar 15 µm. **d** Chronic or repetitive stress do not cause visible changes to TDP-43 subcellular distribution or its nuclear condensation. Representative images for SH-SY5Y cells are shown. Scale bar, 10 µm. **e** Chronic stress leads to a reduction in global protein translation. Translation efficiency was analysed by ribopuromycilation in SH-SY5Y cells after applying the chronic stress paradigm as shown in a. Representative western blot is shown. **f** STMN2 is a short-lived protein, which is more stable in postmitotic as compared to proliferative neuronal cells, as demonstrated by cycloheximide (CHX) pulse-chase. Rocaglamide-A and sylvestrol were included as additional controls – translational inhibitors (30-min treatment). Representative western blots and quantification by densitometry are shown. N=3. **g** Chronic stress causes Golgi apparatus fragmentation in human motor neurons. Representative images and quantification are shown. Cells were analysed at the end of the 48-hour stress protocol as in a. **h** Chronic but not repetitive stress results in increased cell death in SH-SY5Y cultures. Cell viability was measured on Incucyte using a caspase-3/7 dye. Data were normalised to the 0-h time-point. N=3; ****p<0.0001, one-way ANOVA with Tukey’s post-hoc test. **i** Chronic but not repetitive stress results in increased cell death in human motor neuron cultures. Cell viability was measured on Incucyte using a caspase-3/7 dye. Data were normalised to the 0-h time-point. N=4; ****p<0.0001, one-way ANOVA with Tukey’s post-hoc test.

Phenotypically, chronic but not repetitive stress caused Golgi complex disruption in neurons, with STMN2 dissociation and redistribution in the cytosol (Fig. 5g). Longitudinal analysis of cell fitness on Incucyte showed that chronic but not repetitive stress leads to a dramatic decrease in cell viability via apoptosis, both in SH-SY5Y cells and human motor neurons (Fig. 5h,i). Both repetitive and chronic stress impacted cell proliferation in SH-SY5Y cultures, however chronic stress was more detrimental (Fig.S4e).

### Low pre-stress STMN2 sensitises neuronal cells to acute stress

We next asked whether altered baseline levels of STMN2 would affect neuronal fitness and survival during acute stress. We performed survival analysis in SH-SY5Y cells transfected with STMN2-specific or scrambled siRNA. Successful STMN2 knockdown was confirmed by western blot in cultures prepared in parallel (Fig. 6a). Cells were subjected to NaAsO_2_ stress 24 h post-transfection, for 1 hour, and cell proliferation rates and apoptosis rates were monitored by automated imaging on Incucyte for 24 hours of recovery. STMN2 depletion *per se* was well-tolerated by neuroblastoma cells and did not significantly affect their proliferation or viability (Fig. 6b). However, STMN2 siRNA-transfected cells demonstrated a significant increase in cell death and inhibited proliferation post-stress, as compared to the scrambled siRNA control (Fig. 6b).

**Fig. 6.**
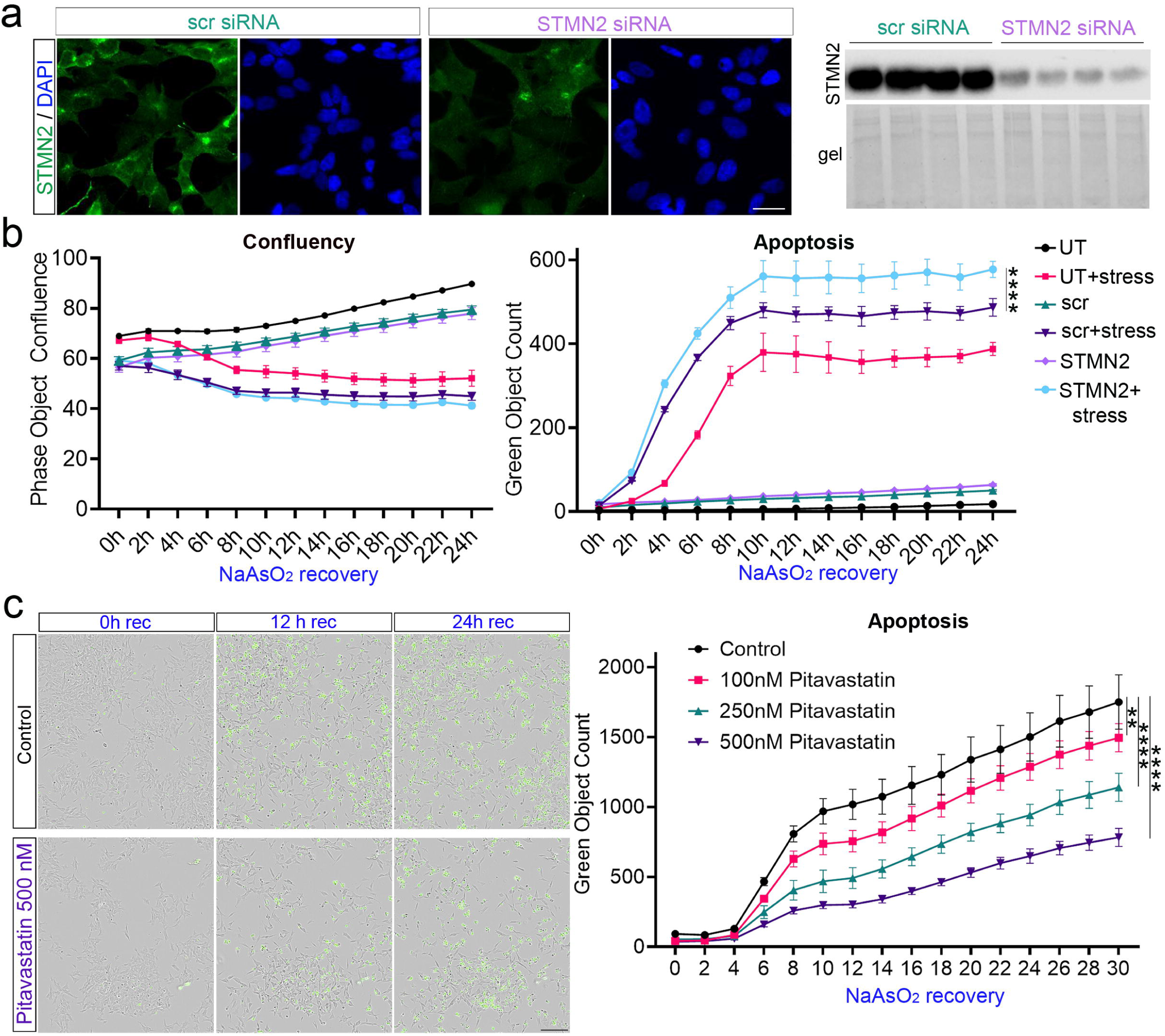
Low pre-stress STMN2 impairs cell survival under stress. **a** Efficient siRNA-mediated STMN2 knockdown. Representative images and western blot are shown. N=4. Cells were analysed 24 h post-transfection. Scale bar 10 µm. **b** STMN2 knockdown does not affect cell proliferation or viability under basal conditions, but is associated with impaired proliferation and survival under stress. Cells were stressed with NaAsO_2_ 24 hours post-transfection, washed and imaged during the recovery period. N=3, ****p<0.0001, one-way ANOVA with Tukey’s post-hoc test. **c** Statin pre-treatment protects neuronal cells from stress-induced apoptosis. Cells were pre-treated with pitavastatin for 16 hours, stressed with NaAsO_2_, washed and imaged during the recovery period. N=3; **p<0.01, ****p<0.0001, one-way ANOVA with Dunnett’s post-hoc test. Scale bar, 100 μm. SH-SY5Y cells were used.

In order to perform a reciprocal experiment, we resorted to pharmacological STMN2 upregulation, using statins. We found that statins at concentrations >500 nM promote neuron-like phenotypes in neuroblastoma cells – most notably, neurite outgrowth (Fig. S1c), which could confound imaging-based quantification. Therefore, lower statin concentrations not associated with visible morphological changes were employed. Cells were pre-treated with pitavastatin for 16 hours, at 100, 250 and 500 nM, and subjected to arsenite stress. Longitudinal analysis on Incucyte revealed a protective effect of pitavastatin pre-treatment during the recovery from NaAsO_2_ stress, compared to the “no-statin” condition, in a concentration-dependent manner (Fig. 6c).

### STMN2 expression compensation at the RNA level in ALS

STMN2 studies in human post-mortem samples have been primarily focused on STMN2 CE-containing RNA so far, with only two reports of STMN2 protein depletion in the ALS spinal cord – one by immunochemistry and another one by single-cell proteomics (1, 47). We tested two commercial anti-STMN2 antibodies, rabbit polyclonal and mouse monoclonal (both Proteintech), which performed well in ICC and recognised a single band in western blot. The rabbit antibody provided significantly stronger staining of spinal motor neurons than the mouse antibody, albeit with some background, including astrocyte and neuropil labelling, but no lipofuscin staining (Fig. 7a, Fig. S5). Given prominent signal in motor neurons, this antibody was deemed suitable for a systematic analysis of STMN2 protein levels in ALS spinal motor neurons. Firstly, a cohort of sporadic ALS (sALS) cases with confirmed TDP-43 pathology was analysed (n=5 per group). Automated analysis of STMN2 pixel intensity in the spinal cord using QuPath demonstrated significant STMN2 downregulation in the surviving motor neurons in sALS as compared to controls (Fig. 7a,b). We were also able to visualise STMN2 mRNA depletion in sALS motor neurons using BaseScope® ISH (Fig. 7c).

**Fig. 7.**
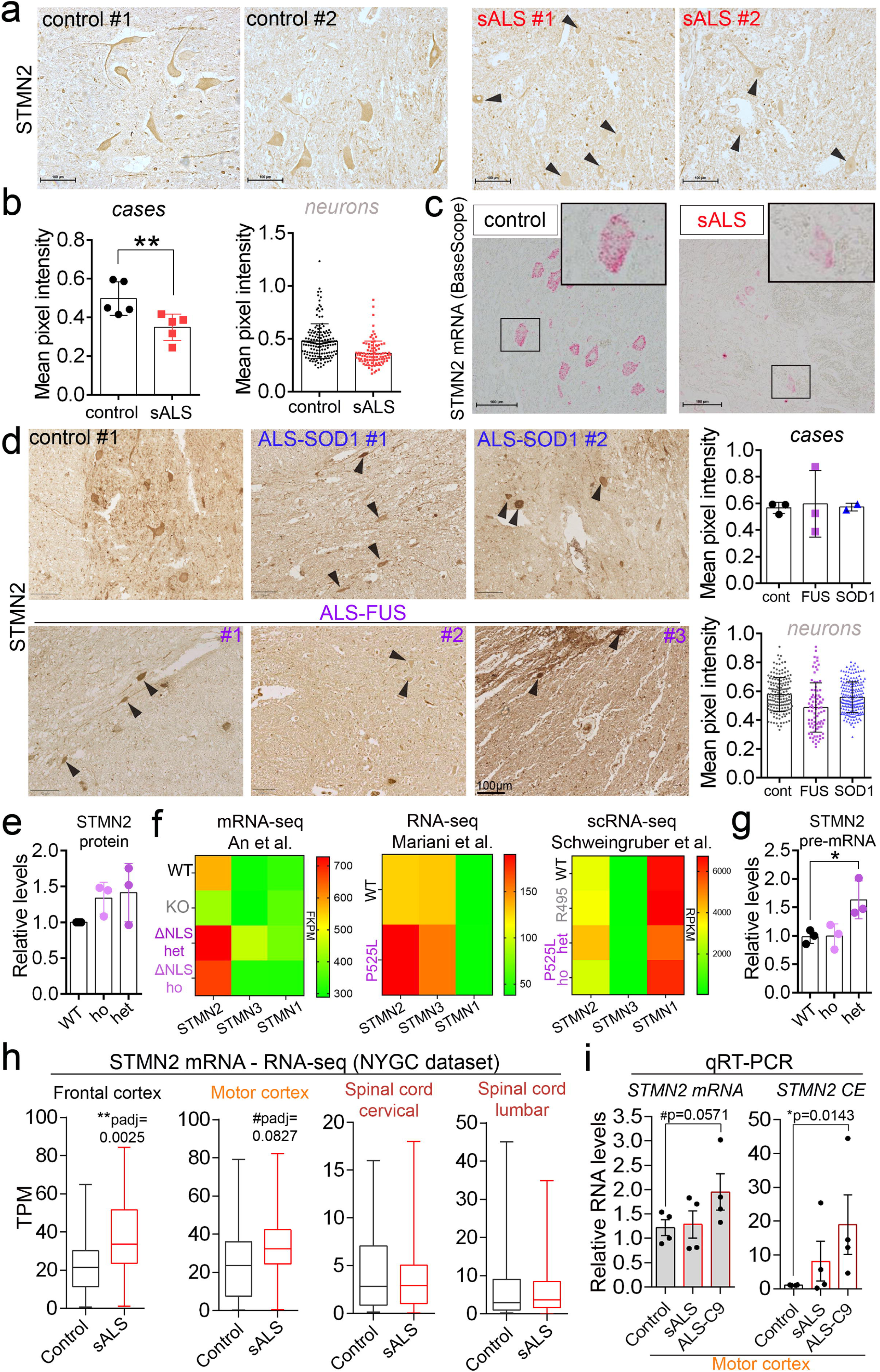
STMN2 expression compensation at the RNA level in ALS. **a,b** STMN2 protein is significantly depleted in spinal motor neurons of sporadic ALS (sALS) patients. Representative images (a) and quantification (b) are shown. N=5; 153 and 96 individual motor neurons analysed for control and sALS, respectively. Arrowheads indicate motor neurons. **p<0.01, Mann-Whitney *U* test. Scale bars, 100 µm. **c** STMN2 mRNA is downregulated in sALS motor neurons. STMN2 full-length mRNA was detected by BaseScope ISH using an exon 4/5-specific probe. Representative images are shown. Scale bars, 100 µm. **d** STMN2 protein level is sustained in non-TDP ALS subtypes. Representative images and quantification are shown. N=3, 2 and 3 for healthy controls, ALS-FUS and ALS-SOD1, respectively; 163, 84 and 208 individual motor neurons analysed for controls, ALS-FUS and ALS-SOD1, respectively. Arrowheads indicate motor neurons. Scale bars, 100 µm. **e** STMN2 protein level is sustained in a physiological ALS-FUS cell model. Quantification for FUSΔNLS SH-SY5Y lines is shown. N=3. **f** STMN2 mRNA upregulation in ALS-FUS cell models. **g**, STMN2 pre-mRNA upregulation in a physiological ALS-FUS cell model. Quantification for FUSΔNLS SH-SY5Y lines is shown. *p=0.05, Mann-Whitney *U* test. N=3. **g** STMN2 mRNA levels in the CNS area of sALS patients with low vs. high pathology burden. Bulk RNA-seq datasets from the NYGC ALS consortium were used (∼200 sALS and ALS-free patients). **h** STMN2 RNA analysis in ALS-TDP motor cortex using qRT-PCR. N=4 per group, one-tailed Mann-Whitney *U* test.

Since our cellular findings suggested that STMN2 can be misregulated via mechanisms other than TDP-43 loss of function, we performed STMN2 protein analysis in a cohort of non-TDP ALS cases – those caused by *SOD1* or *FUS* mutations. Both ALS subtypes are characterised by altered translation and/or SG-related pathology (48–54). Two ALS-SOD1 and three ALS-FUS cases were included in the study (55, 56). One of the ALS-FUS cases was a new, unpublished case (p.Pro525Arg) with abundant FUS pathology (Darwin et al., submitted). Although spinal neurons in these non-TDP ALS cases tended to have variable STMN2 protein expression, on average, they showed no significant depletion, as compared to controls (Fig. 7d).

Consistent with the ALS-FUS tissue data, STMN2 protein level was maintained in FUSΔNLS cell lines, both under basal conditions and under stress (Fig. 7e; Fig.S6a), despite a stress-mimicking state in these lines (i.e. phospho-eIF2α upregulation) (Fig.S6b). This led us to hypothesise that in non-TDP ALS, the defects in STMN2 protein production due to translation/SG misregulation may be compensated at the RNA level. To test this, we re-analysed published RNA-seq data from ALS-FUS cell models. Indeed, STMN2 mRNA levels were found upregulated in two neuroblastoma cell models – FUSΔNLS lines (both homo- and heterozygous) (26) and in a cell line with inducible expression of the P525L variant (57) (Fig. 7f). STMN2 mRNA was also found upregulated in iPSC-derived motor neurons with heterozygous expression of the P525L mutant (single-cell RNA-seq) (58) (Fig. 7f). In contrast, other stathmins (STMN1 and -3) did not show differential expression in ALS-FUS models. Consistently, we also detected STMN2 pre-mRNA upregulation in FUSΔNLS cells (Fig. 7g).

TDP-43 pathology can lead to translation and SG defects (59–62), therefore an RNA-level compensation may also occur in this subtype at the early disease stages, before the onset of TDP-43 proteinopathy. We performed STMN2 mRNA analysis in human post-mortem tissue from sALS patients and healthy controls using bulk RNA-seq datasets from the New York Genome Center (NYGC) ALS Consortium (n=∼200 for patients and controls in total). CNS regions with different extent of neuropathology (motor cortex, frontal cortex and spinal cord) were included in this analysis. STMN2 mRNA was found significantly upregulated in the frontal cortex of ALS patients (padj=0.0025), with a trend towards STMN2 upregulation in the motor cortex (padj=0.08227) – the two CNS regions relatively spared in the disease, whereas no difference was observed for the spinal cord (Fig. 7h). Consistently, we also detected a trend towards STMN2 mRNA upregulation in the motor cortex in a small ALS-C9 cohort (n=4) using qRT-PCR (Fig. 7h). Interestingly, ALS-C9 cases in this latter sample set were characterised by more dramatic STMN2 CE accumulation than sALS cases (Fig. 7h). Finally, we found a trend towards STMN2 mRNA upregulation in the frontal cortex of ALS-SOD1 patients in the NYGC ALS cohort (Fig.S7), although this analysis was limited by the small number of SOD1 cases.

## DISCUSSION

Our detailed analysis of STMN2 regulation under acute and chronic stress reveals how intrinsic sensitivity of STMN2 translation to stress-induced molecular changes creates a vulnerable neuronal state (Fig. 8). Our findings implicate two common molecular denominators in ALS – misregulation of translation and SGs – in STMN2 downregulation, independently of TDP-43 loss of function in splicing (Fig. 8).

**Fig. 8.**
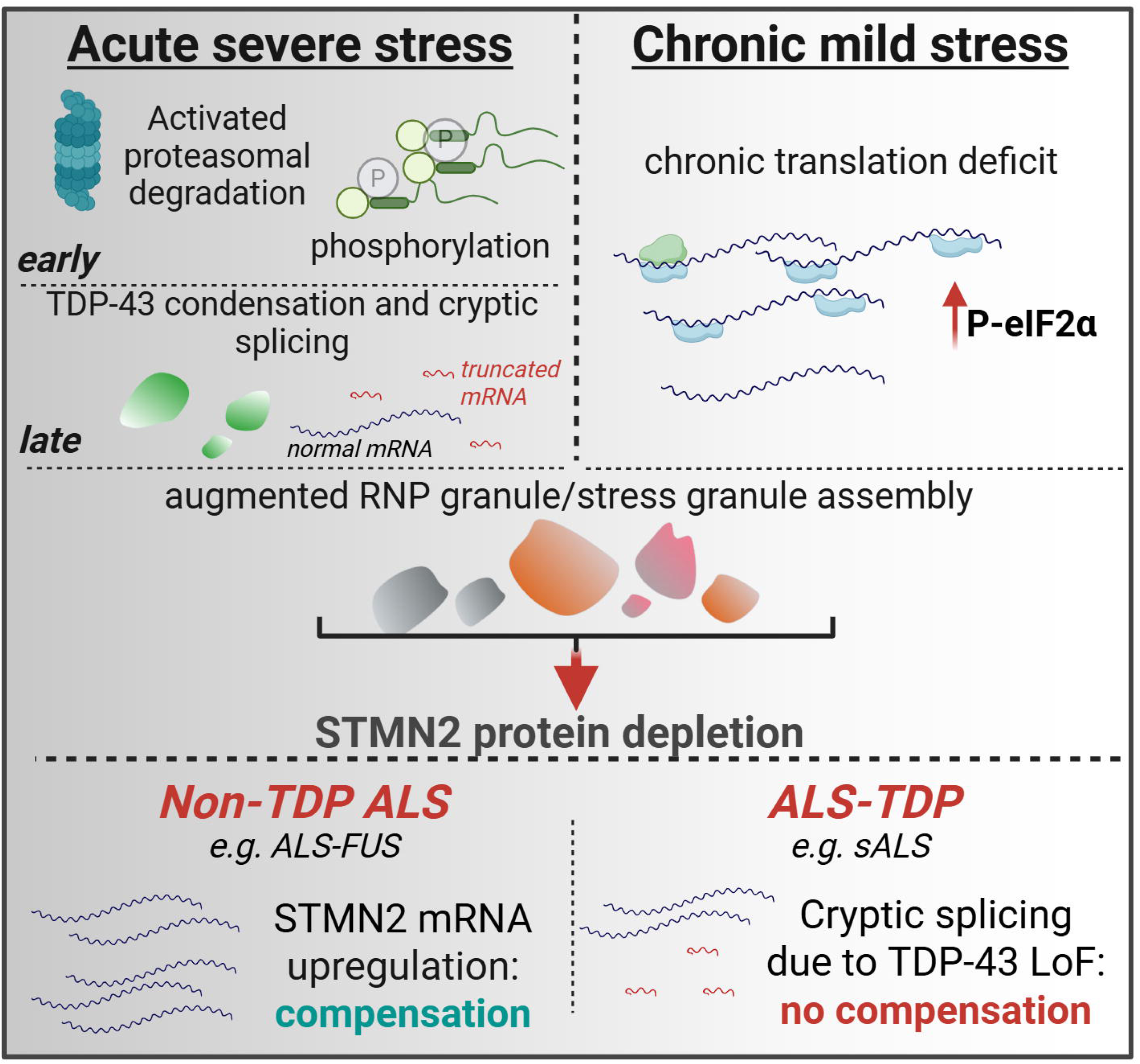
Model for STMN2 dysregulation and compensation in ALS.

Here, we describe the “early” and “late” mechanisms of STMN2 depletion during stress. Firstly, we confirm the causal effect of TDP-43 condensation on STMN2 depletion late in stress/recovery in direct experiments, using murine cells lacking the cryptic exon-dependent regulatory mechanism. Secondly, we find that STMN2 depletion/inactivation in acute stress can occur almost immediately after the exposure to stressor and independently of cryptic splicing – via activated proteasomal degradation and SG-mediated translation repression. Enhanced STMN2 degradation via the proteasome may represent a side-effect of stress-induced proteasome activation – required to prevent the build-up of misfolded proteins during stress (63). Likewise, STMN2 translation suppression by SGs may be secondary to their role in selective translation critical for proteostasis maintenance under stress (39). Thus, these two cytoprotective mechanisms may lead to “unintentional”, non-specific STMN2 reduction during stress. In contrast, STMN2 phosphorylation was identified as a targeted, physiological mechanism of its functional inactivation, through prevention of tubulin binding (32, 64). However, a recent study has suggested that tubulin binding by STMN2 is dispensable for axonal regeneration (65). Therefore, phosphorylation under stress may not be linked to *regulated* loss of STMN2 activity, instead representing a side-effect of increased kinase activity. On the other hand, we show that STMN2 translation can withstand global stress-induced translation shutdown, remaining high during stress (∼30% of basal). Therefore, post-transcriptional (TDP-43-regulated cryptic splicing and SGs) and post-translational (phosphorylation) regulation of STMN2 levels/activity may represent physiological mechanisms – presumably more controllable and fail-safe than regulation of the translational output of the *STMN2* gene. Further studies are required to establish to what extent stress-induced STMN2 depletion is a controlled event with a physiological purpose and to what extent – an unwanted and potentially deleterious consequence of the integrated stress response activation.

We find that STMN2 expression is also significantly impacted by chronic translation deficits. Consistently, in a recent study using machine learning analysis of proteomic datasets, a decrease in ribosomal protein RPS29 was a shared feature of ALS cell models, which was associated with STMN2 downregulation (66). STMN2’s sensitivity to prolonged reduction in translation is defined by its extremely rapid turnover. Our data on STMN2 half-life in motor neurons (∼1.5-2 hours) are consistent with the previous studies in neurons (29); however, we find that its turnover is more rapid in proliferating cells (half-life of ∼50 min). Short-lived proteins, defined as those with a half-live ≤8 hours, account for only ∼5% of the human proteome, and most of them are low-abundance, regulatory proteins involved in signalling and cell cycle regulation (67). Although this latter study did not use neuronal cells, for which the short-lived protein repertoire may be different, STMN2 is among the most labile proteins. This property enables its efficient control but at the same time increases vulnerability to pathological states with impaired translation.

Based on our data, we speculate that the magnitude and duration of stress dictate the level of STMN2 expression impairment. It is possible that severe acute stresses causing TDP-43 nuclear condensation and STMN2 protein clearance are irreversible, causing irreparable damage eventually leading to neuronal death. It cannot be excluded that neurons in the human body sometimes experience such stress. However, low-grade chronic stress associated with mild but prolonged translational disturbances is a more likely scenario in the disease context. Our data suggests that STMN2 downregulation creates a vulnerable neuronal state – not only related to axonal health but also at the whole cell level.

Misregulation of SGs, a well-known feature of stress response, has been reported across several neurodegenerative diseases (68). Although SG assembly is typical for acute stress response (69), chronic stress states may also be associated with an increase in submicroscopic RNP granulation/condensation – consistent with the coalescence of pre-existing RNP complexes under stress (70, 71). SG regulation is very different in cultured cells and *in vivo* as well as in neuronal *vs.* non-neuronal cells. Indeed, in contrast to cell culture studies, SG induction has only been achieved in a handful of mammalian *in vivo* models, requiring the use of harsh, terminal stresses (23, 72, 73). Neurons also appear to be more resistant to large SG assembly (23, 74). Submicroscopic RNP condensation may play a more important role in STMN2 (mis)regulation via translational repression in disease than the assembly of large SGs (Fig. 8). Some recent studies using non-neuronal cells questioned the SG role in global translation repression (41, 75). Our data indicate that SG function in translation is transcript- and cell type-specific, where the highly abundant and actively translated neurospecific STMN2 mRNA is sensitive to SG formation. Interestingly, in our studies, we noticed that endogenous STMN2 protein was consistently enriched in cytoplasmic foci resembling SGs, which were confirmed to correspond to SGs using anti-G3BP1 co-staining (Fig.S8). However, we were unable to confirm STMN2 enrichment in SGs using ectopically expressed GFP-tagged STMN2 (Fig S8). Anti-STMN2 antibodies may non-specifically recognise SG components due to their high density within SGs. Yet we cannot rule out that STMN2 is indeed a SG component, where a tag on its N-terminus impairs its SG recruitment. Further studies into potential protein-level regulation of STMN2 by SGs would be required.

Translation restoration using small molecules has long been considered an attractive therapeutic strategy in neurodegenerative diseases, including ALS (76). Some of such translation activators lead to SG dissolution (77). For example, DNL343, an investigational small-molecule drug – translation activator with a mechanism of action similar to ISRIB – showed promise in reversing pathological phenotypes in TDP-43 transgenic mice (78). Other compounds, with less favourable pharmacokinetic profiles, also showed promise in earlier reports (79, 80). Studies into the effect of these compounds specifically on STMN2 regulation in human neuronal cell models are warranted, across different ALS subtypes, to establish whether their beneficial effect is at least partially mediated by STMN2 maintenance/restoration.

Our cellular and human tissue data suggest that STMN2 is subject to expression compensation at the RNA level in ALS – which may be sufficient to counteract the translation deficits in non-TDP ALS such as ALS-FUS (Fig.8). We find evidence for such compensation in the CNS regions in ALS-TDP that are relatively spared (cortex), but not in those with profound TDP-43 pathology (spinal cord). In the spinal cord which is severely affected, this compensation likely failed due to cryptic splicing. Since TDP-43 itself affects SGs (62) and translation regulation (81, 82), its pathology would create a vicious cycle of cellular dysfunction. Interestingly, STMN2 mRNA upregulation was reported in two independent ALS-SOD1 mouse models (5, 83). However, we found STMN2 is differently regulated in human and mouse cells, including under stress – being resistant to stress-induced clearance in mouse cells. STMN2 preservation in ALS-FUS and -SOD1 may ameliorate the disease course in these ALS subtypes. Yet, pathological changes in these subtypes still develop and progress, despite STMN2 preservation. This suggests that multi-target interventions may be required to achieve a significant therapeutic effect in ALS.

In conclusion, we demonstrate that molecular pathologies other than TDP-43 dysfunction converge on STMN2 and provide an additional mechanism for neuronal impairment by abnormal translation and SG regulation. While confirming vulnerability of neurons with developing TDP-43 dysfunction, our study supports the development of therapies restoring translation and RNP granule regulation in ALS subtypes with and without TDP-43 pathology.

## Acknowledgements

We are grateful to the Sheffield Brain Tissue Bank for supplying the tissue and to the NYGC ALS consortium for providing datasets for analysis. We thank those who have donated tissue for scientific research and their families who have supported this. We also thank Dan Fillingham for assisting with human tissue preparation. We are grateful to Ze’ev Melamed, Guillaume Hautbergue and Alison Twelvetrees for useful discussions.

## Funding

The work was supported by the UKRI Future Leaders Fellowship (MR/W004615/1), responsive mode grants from the MRC (MR/W028522/1) and BBSRC (BB/V014110/1), MND Association fellowship/grant (968–799), and Alzheimer’s Research UK pilot grant (ARUK-PPG2023B-007) to T.A.S. B.S.C.E. was supported by a PhD studentship from the MND Scotland (2022/MNDS/PHD/8045SHEL; to T.A.S. and K.J.D.V.). V.K. is funded by a PhD studentship from the University of Sheffield Faculty of Health (to T.A.S.). J.C.-K. is supported by a Lister Institute Research Prize and Tambourine’s ALS Breakthrough Research Fund in partnership with the Milken Institute Center for Strategic Philanthropy. J.R.H. acknowledges funding from The Pathological Society of Great Britain and Ireland and the MND Association. M.N. was supported by the Association for Frontotemporal Degeneration Holloway Postdoctoral Fellowship, The Cullen Education and Research Foundation, and Hop On a Cure.

## Author contributions

T.A.S. supervised the study, designed and conducted experiments, analysed data and wrote the manuscript. B.C.S.E. designed and conducted experiments, analysed data and wrote the manuscript. A.S.A., W.-P.H., S.J.J., S.B., R.E.H. and V.K. designed and conducted experiments and analysed data. M.N. and C.L.-T. contributed study tools. S.G.C., K.J.D.V., J.R.H. and J.C.-K. contributed to the study design, data analysis or study supervision. All authors contributed to the manuscript writing and approved its final version.

## Data and materials availability

Data and materials generated in this study will be made available by the corresponding author, upon reasonable request, subject to a materials transfer agreement.

## Conflicts of interest

C.L.T. and M.N. have filed patent applications related to the use of statins in neurodegenerative diseases. Other authors declare no competing interests.

## Notes

### Competing Interest Statement

CLT and MN have filed patent applications related to the use of statins in neurodegenerative diseases. Other authors declare no competing interests.

## References

1 Klim, J.R., Williams, L.A., Limone, F., Guerra San Juan, I., Davis-Dusenbery, B.N., Mordes, D.A., Burberry, A., Steinbaugh, M.J., Gamage, K.K., Kirchner, R., et al. (2019) ALS-implicated protein TDP-43 sustains levels of STMN2, a mediator of motor neuron growth and repair. Nat Neurosci, 22, 167–179.

2 Melamed, Z., Lopez-Erauskin, J., Baughn, M.W., Zhang, O., Drenner, K., Sun, Y., Freyermuth, F., McMahon, M.A., Beccari, M.S., Artates, J.W. et al. (2019) Premature polyadenylation-mediated loss of stathmin-2 is a hallmark of TDP-43-dependent neurodegeneration. Nat Neurosci, 22, 180–190.

3 Agra Almeida Quadros, A.R., Li, Z., Wang, X., Ndayambaje, I.S., Aryal, S., Ramesh, N., Nolan, M., Jayakumar, R., Han, Y., Stillman, H., et al. (2024) Cryptic splicing of stathmin-2 and UNC13A mRNAs is a pathological hallmark of TDP-43-associated Alzheimer’s disease. Acta Neuropathol, 147, 9.

4 Baughn, M.W., Melamed, Z., Lopez-Erauskin, J., Beccari, M.S., Ling, K., Zuberi, A., Presa, M., Gonzalo-Gil, E., Maimon, R., Vazquez-Sanchez, S. et al. (2023) Mechanism of STMN2 cryptic splice-polyadenylation and its correction for TDP-43 proteinopathies. Science, 379, 1140–1149.

5 Bellouze, S., Baillat, G., Buttigieg, D., de la Grange, P., Rabouille, C. and Haase, G. (2016) Stathmin 1/2-triggered microtubule loss mediates Golgi fragmentation in mutant SOD1 motor neurons. Mol Neurodegener, 11, 43.

6 Wang, Q., Zhang, Y., Wang, M., Song, W.M., Shen, Q., McKenzie, A., Choi, I., Zhou, X., Pan, P.Y., Yue, Z. et al. (2019) The landscape of multiscale transcriptomic networks and key regulators in Parkinson’s disease. Nat Commun, 10, 5234.

7 San Juan, I.G., Nash, L.A., Smith, K.S., Leyton-Jaimes, M.F., Qian, M., Klim, J.R., Limone, F., Dorr, A.B., Couto, A., Pintacuda, G., et al. (2022) Loss of mouse Stmn2 function causes motor neuropathy. Neuron, 110, 4031.

8 Krus, K.L., Strickland, A., Yamada, Y., Devault, L., Schmidt, R.E., Bloom, A.J., Milbrandt, J. and DiAntonio, A. (2022) Loss of Stathmin-2, a hallmark of TDP-43-associated ALS, causes motor neuropathy. Cell Rep, 39, 111001.

9 Liedtke, W., Leman, E.E., Fyffe, R.E., Raine, C.S. and Schubart, U.K. (2002) Stathmin-deficient mice develop an age-dependent axonopathy of the central and peripheral nervous systems. Am J Pathol, 160, 469–480.

10 Zhu, X.X., Kozarsky, K., Strahler, J.R., Eckerskorn, C., Lottspeich, F., Melhem, R., Lowe, J., Fox, D.A., Hanash, S.M. and Atweh, G.F. (1989) Molecular cloning of a novel human leukemia-associated gene. Evidence of conservation in animal species. J Biol Chem, 264, 14556–14560.

11 Sun, S., Sun, Y., Ling, S.C., Ferraiuolo, L., McAlonis-Downes, M., Zou, Y., Drenner, K., Wang, Y., Ditsworth, D., Tokunaga, S. et al. (2015) Translational profiling identifies a cascade of damage initiated in motor neurons and spreading to glia in mutant SOD1-mediated ALS. Proc Natl Acad Sci U S A, 112, E6993–7002.

12 Sobel, A., Boutterin, M.C., Beretta, L., Chneiweiss, H., Doye, V. and Peyro-Saint-Paul, H. (1989) Intracellular substrates for extracellular signaling. Characterization of a ubiquitous, neuron-enriched phosphoprotein (stathmin). J Biol Chem, 264, 3765–3772.

13 Shin, J.E., Geisler, S. and DiAntonio, A. (2014) Dynamic regulation of SCG10 in regenerating axons after injury. Exp Neurol, 252, 1–11.

14 Morii, H., Shiraishi-Yamaguchi, Y. and Mori, N. (2006) SCG10, a microtubule destabilizing factor, stimulates the neurite outgrowth by modulating microtubule dynamics in rat hippocampal primary cultured neurons. J Neurobiol, 66, 1101–1114.

15 Huang, W.P., Ellis, B.C.S., Hodgson, R.E., Sanchez Avila, A., Kumar, V., Rayment, J., Moll, T. and Shelkovnikova, T.A. (2024) Stress-induced TDP-43 nuclear condensation causes splicing loss of function and STMN2 depletion. Cell Rep, 43, 114421.

16 Al-Chalabi, A., Calvo, A., Chio, A., Colville, S., Ellis, C.M., Hardiman, O., Heverin, M., Howard, R.S., Huisman, M.H.B., Keren, N. et al. (2014) Analysis of amyotrophic lateral sclerosis as a multistep process: a population-based modelling study. Lancet Neurol, 13, 1108–1113.

17 Qureshi, D., Grubic, N., Maxwell, C.J., Bush, S.H., Casey, G., Isenberg, S.R., Tanuseputro, P. and Webber, C. (2024) Association of Disease Trajectory and Place of Care with End-of-Life Burdensome Transitions: A Retrospective Cohort Study. J Am Med Dir Assoc, 25, 105229.

18 Chapman, L., Cooper-Knock, J. and Shaw, P.J. (2023) Physical activity as an exogenous risk factor for amyotrophic lateral sclerosis: a review of the evidence. Brain, 146, 1745–1757.

19 Shelkovnikova, T.A., An, H., Skelt, L., Tregoning, J.S., Humphreys, I.R. and Buchman, V.L. (2019) Antiviral Immune Response as a Trigger of FUS Proteinopathy in Amyotrophic Lateral Sclerosis. Cell Rep, 29, 4496–4508 e4494.

20 Bellmann, J., Monette, A., Tripathy, V., Sojka, A., Abo-Rady, M., Janosh, A., Bhatnagar, R., Bickle, M., Mouland, A.J. and Sterneckert, J. (2019) Viral Infections Exacerbate FUS-ALS Phenotypes in iPSC-Derived Spinal Neurons in a Virus Species-Specific Manner. Front Cell Neurosci, 13, 480.

21 Yang, J., Li, Y., Wang, S., Li, H., Zhang, L., Zhang, H., Wang, P.H., Zheng, X., Yu, X.F. and Wei, W. (2023) The SARS-CoV-2 main protease induces neurotoxic TDP-43 cleavage and aggregates. Signal Transduct Target Ther, 8, 109.

22 Hipp, M.S., Kasturi, P. and Hartl, F.U. (2019) The proteostasis network and its decline in ageing. Nat Rev Mol Cell Biol, 20, 421–435.

23 Shelkovnikova, T.A., Dimasi, P., Kukharsky, M.S., An, H., Quintiero, A., Schirmer, C., Buee, L., Galas, M.C. and Buchman, V.L. (2017) Chronically stressed or stress-preconditioned neurons fail to maintain stress granule assembly. Cell Death Dis, 8, e2788.

24 Shelkovnikova, T.A., Kukharsky, M.S., An, H., Dimasi, P., Alexeeva, S., Shabir, O., Heath, P.R. and Buchman, V.L. (2018) Protective paraspeckle hyper-assembly downstream of TDP-43 loss of function in amyotrophic lateral sclerosis. Mol Neurodegener, 13, 30.

25 An, H., Elvers, K.T., Gillespie, J.A., Jones, K., Atack, J.R., Grubisha, O. and Shelkovnikova, T.A. (2022) A toolkit for the identification of NEAT1_2/paraspeckle modulators. Nucleic Acids Res, 50, e119.

26 An, H., Skelt, L., Notaro, A., Highley, J.R., Fox, A.H., La Bella, V., Buchman, V.L. and Shelkovnikova, T.A. (2019) ALS-linked FUS mutations confer loss and gain of function in the nucleus by promoting excessive formation of dysfunctional paraspeckles. Acta Neuropathol Commun, 7, 7.

27 Ritchie, M.E., Phipson, B., Wu, D., Hu, Y., Law, C.W., Shi, W. and Smyth, G.K. (2015) limma powers differential expression analyses for RNA-sequencing and microarray studies. Nucleic Acids Res, 43, e47.

28 Humphrey, J., Venkatesh, S., Hasan, R., Herb, J.T., de Paiva Lopes, K., Kucukali, F., Byrska-Bishop, M., Evani, U.S., Narzisi, G., Fagegaltier, D., et al. (2023) Integrative transcriptomic analysis of the amyotrophic lateral sclerosis spinal cord implicates glial activation and suggests new risk genes. Nat Neurosci, 26, 150–162.

29 Deng, X., Bradshaw, G., Kalocsay, M. and Mitchison, T. (2025) Tubulin Regulates the Stability and Localization of STMN2 by Binding Preferentially to Its Soluble Form. bioRxiv, doi: 10.1101/2025.02.27.640326.

30 Berkers, C.R., van Leeuwen, F.W., Groothuis, T.A., Peperzak, V., van Tilburg, E.W., Borst, J., Neefjes, J.J. and Ovaa, H. (2007) Profiling proteasome activity in tissue with fluorescent probes. Mol Pharm, 4, 739–748.

31 Dar, S.A., Malla, S., Martinek, V., Payea, M.J., Lee, C.T., Martin, J., Khandeshi, A.J., Martindale, J.L., Belair, C. and Maragkakis, M. (2024) Full-length direct RNA sequencing uncovers stress-granule dependent RNA decay upon cellular stress. eLife, 10.7554/eLife.96284.2

32 Steinmetz, M.O., Jahnke, W., Towbin, H., Garcia-Echeverria, C., Voshol, H., Muller, D. and van Oostrum, J. (2001) Phosphorylation disrupts the central helix in Op18/stathmin and suppresses binding to tubulin. EMBO Rep, 2, 505–510.

33 Tararuk, T., Ostman, N., Li, W., Bjorkblom, B., Padzik, A., Zdrojewska, J., Hongisto, V., Herdegen, T., Konopka, W., Courtney, M.J. et al. (2006) JNK1 phosphorylation of SCG10 determines microtubule dynamics and axodendritic length. J Cell Biol, 173, 265–277.

34 Di Paolo, G., Lutjens, R., Pellier, V., Stimpson, S.A., Beuchat, M.H., Catsicas, S. and Grenningloh, G. (1997) Targeting of SCG10 to the area of the Golgi complex is mediated by its NH2-terminal region. J Biol Chem 272, 5175–5182.

35 tom Dieck, S., Kochen, L., Hanus, C., Heumuller, M., Bartnik, I., Nassim-Assir, B., Merk, K., Mosler, T., Garg, S., Bunse, S., et al. (2015) Direct visualization of newly synthesized target proteins in situ. Nat Methods, 12, 411–414.

36 Tian, A.L., Wu, Q., Liu, P., Zhao, L., Martins, I., Kepp, O., Leduc, M. and Kroemer, G. (2021) Lysosomotropic agents including azithromycin, chloroquine and hydroxychloroquine activate the integrated stress response. Cell Death Dis, 12, 6.

37 Naineni, S.K., Bhatt, G., Jiramongkolsiri, E., Robert, F., Cencic, R., Huang, S., Nagar, B. and Pelletier, J. (2024) Protein-RNA interactions mediated by silvestrol-insight into a unique molecular clamp. Nucleic Acids Res, 52, 12701–12711.

38 Hofmann, S., Kedersha, N., Anderson, P. and Ivanov, P. (2021) Molecular mechanisms of stress granule assembly and disassembly. Biochim Biophys Acta Mol Cell Res, 1868, 118876.

39 Buchan, J.R. and Parker, R. (2009) Eukaryotic stress granules: the ins and outs of translation. Molecular cell, 36, 932–941.

40 Baymiller, M. and Moon, S.L. (2023) Stress Granules as Causes and Consequences of Translation Suppression. Antioxid Redox Signal, 39, 390–409.

41 Mateju, D., Eichenberger, B., Voigt, F., Eglinger, J., Roth, G. and Chao, J.A. (2020) Single-Molecule Imaging Reveals Translation of mRNAs Localized to Stress Granules. Cell, 183, 1801–1812 e1813.

42 Sidrauski, C., Acosta-Alvear, D., Khoutorsky, A., Vedantham, P., Hearn, B.R., Li, H., Gamache, K., Gallagher, C.M., Ang, K.K., Wilson, C. et al. (2013) Pharmacological brake-release of mRNA translation enhances cognitive memory. Elife, 2, e00498.

43 Uechi, H., Sridharan, S., Nijssen, J., Bilstein, J., Iglesias-Artola, J.M., Kishigami, S., Casablancas-Antras, V., Poser, I., Martinez, E.J., Boczek, E. et al. (2025) Small-molecule dissolution of stress granules by redox modulation benefits ALS models. Nat Chem Biol, 21, 1577–1588

44 Zyryanova, A.F., Kashiwagi, K., Rato, C., Harding, H.P., Crespillo-Casado, A., Perera, L.A., Sakamoto, A., Nishimoto, M., Yonemochi, M., Shirouzu, M. et al. (2021) ISRIB Blunts the Integrated Stress Response by Allosterically Antagonising the Inhibitory Effect of Phosphorylated eIF2 on eIF2B. Molecular cell, 81, 88–103 e106.

45 Lehmkuhl, E.M. and Zarnescu, D.C. (2018) Lost in Translation: Evidence for Protein Synthesis Deficits in ALS/FTD and Related Neurodegenerative Diseases. Adv Neurobiol, 20, 283–301.

46 Luh, L.M. and Bertolotti, A. (2020) Potential benefit of manipulating protein quality control systems in neurodegenerative diseases. Curr Opin Neurobiol, 61, 125–132.

47 Guise, A.J., Misal, S.A., Carson, R., Chu, J.H., Boekweg, H., Van Der Watt, D., Welsh, N.C., Truong, T., Liang, Y., Xu, S., et al. (2024) TDP-43-stratified single-cell proteomics of postmortem human spinal motor neurons reveals protein dynamics in amyotrophic lateral sclerosis. Cell Rep, 43, 113636.

48 Gal, J., Kuang, L., Barnett, K.R., Zhu, B.Z., Shissler, S.C., Korotkov, K.V., Hayward, L.J., Kasarskis, E.J. and Zhu, H. (2016) ALS mutant SOD1 interacts with G3BP1 and affects stress granule dynamics. Acta neuropathologica, 132, 563–576.

49 Dormann, D., Madl, T., Valori, C.F., Bentmann, E., Tahirovic, S., Abou-Ajram, C., Kremmer, E., Ansorge, O., Mackenzie, I.R., Neumann, M. et al. (2012) Arginine methylation next to the PY-NLS modulates Transportin binding and nuclear import of FUS. EMBO J, 31, 4258–4275.

50 Mateju, D., Franzmann, T.M., Patel, A., Kopach, A., Boczek, E.E., Maharana, S., Lee, H.O., Carra, S., Hyman, A.A. and Alberti, S. (2017) An aberrant phase transition of stress granules triggered by misfolded protein and prevented by chaperone function. EMBO J, 36, 1669–1687.

51 Shelkovnikova, T.A., Robinson, H.K., Southcombe, J.A., Ninkina, N. and Buchman, V.L. (2014) Multistep process of FUS aggregation in the cell cytoplasm involves RNA-dependent and RNA-independent mechanisms. Human molecular genetics, 23, 5211–5226.

52 Birsa, N., Ule, A.M., Garone, M.G., Tsang, B., Mattedi, F., Chong, P.A., Humphrey, J., Jarvis, S., Pisiren, M., Wilkins, O.G. et al. (2021) FUS-ALS mutants alter FMRP phase separation equilibrium and impair protein translation. Sci Adv, 7, eabf8660.

53 Kamelgarn, M., Chen, J., Kuang, L., Jin, H., Kasarskis, E.J. and Zhu, H. (2018) ALS mutations of FUS suppress protein translation and disrupt the regulation of nonsense-mediated decay. Proc Natl Acad Sci U S A, 115, E11904–E11913.

54 Szewczyk, B., Gunther, R., Japtok, J., Frech, M.J., Naumann, M., Lee, H.O. and Hermann, A. (2023) FUS ALS neurons activate major stress pathways and reduce translation as an early protective mechanism against neurodegeneration. Cell Rep, 42, 112025.

55 Hewitt, C., Kirby, J., Highley, J.R., Hartley, J.A., Hibberd, R., Hollinger, H.C., Williams, T.L., Ince, P.G., McDermott, C.J. and Shaw, P.J. (2010) Novel FUS/TLS mutations and pathology in familial and sporadic amyotrophic lateral sclerosis. Arch Neurol, 67, 455–461.

56 Shaw, P.J., Tomkins, J., Slade, J.Y., Usher, P., Curtis, A., Bushby, K. and Ince, P.G. (1997) CNS tissue Cu/Zn superoxide dismutase (SOD1) mutations in motor neurone disease (MND). Neuroreport, 8, 3923–3927.

57 Mariani, D., Setti, A., Castagnetti, F., Vitiello, E., Stufera Mecarelli, L., Di Timoteo, G., Giuliani, A., D’Angelo, A., Santini, T., Perego, E., et al. (2024) ALS-associated FUS mutation reshapes the RNA and protein composition of stress granules. Nucleic Acids Res, 52, 13269–13289.

58 Schweingruber, C., Nijssen, J., Mechtersheimer, J., Reber, S., Leboeuf, M., O’Brien, N.L., Mei, I., Hedges, E., Keuper, M., Benitez, J.A. et al. (2025) Single-cell RNA-sequencing reveals early mitochondrial dysfunction unique to motor neurons shared across FUS- and TARDBP-ALS. Nat Commun, 16, 4633.

59 Chu, J.F., Majumder, P., Chatterjee, B., Huang, S.L. and Shen, C.J. (2019) TDP-43 Regulates Coupled Dendritic mRNA Transport-Translation Processes in Co-operation with FMRP and Staufen1. Cell Rep, 29, 3118–3133 e3116.

60 Piol, D., Robberechts, T. and Da Cruz, S. (2023) Lost in local translation: TDP-43 and FUS in axonal/neuromuscular junction maintenance and dysregulation in amyotrophic lateral sclerosis. Neuron, 111, 1355–1380.

61 Blokhuis, A.M., Koppers, M., Groen, E.J.N., van den Heuvel, D.M.A., Dini Modigliani, S., Anink, J.J., Fumoto, K., van Diggelen, F., Snelting, A., Sodaar, P., et al. (2016) Comparative interactomics analysis of different ALS-associated proteins identifies converging molecular pathways. Acta neuropathologica, 132, 175–196.

62 Aulas, A. and Vande Velde, C. (2015) Alterations in stress granule dynamics driven by TDP-43 and FUS: a link to pathological inclusions in ALS? Front Cell Neurosci, 9, 423.

63 Lee, D. and Goldberg, A.L. (2022) 26S proteasomes become stably activated upon heat shock when ubiquitination and protein degradation increase. Proc Natl Acad Sci U S A, 119, e2122482119.

64 Wittmann, T., Bokoch, G.M. and Waterman-Storer, C.M. (2004) Regulation of microtubule destabilizing activity of Op18/stathmin downstream of Rac1. J Biol Chem, 279, 6196–6203.

65 Beccari, M.S., Arnold-Garcia, O., Baughn, M.W., Artates, J.W., McAlonis-Downes, M., Lim, J., Leyva-Cazares, D.F., Rubio-Lara, H.I., Ramirez-Rodriguez, A., Bernal-Buenrostro, C.N. et al. (2025) Stathmin-2 enhances motor axon regeneration after injury independent of its binding to tubulin. Proc Natl Acad Sci U S A, 122, e2502294122.

66 Xu, W., Guo, Z., Guan, Y., Lv, S., Gao, X., Luo, W., Cheng, T., Shao, Z., Tao, B., Wang, T. et al. (2025) Machine learning-based proteomics profiling of ALS identifies downregulation of RPS29 that maintains protein homeostasis and STMN2 level. Commun Biol, 8, 1177.

67 Li, J., Cai, Z., Vaites, L.P., Shen, N., Mitchell, D.C., Huttlin, E.L., Paulo, J.A., Harry, B.L. and Gygi, S.P. (2021) Proteome-wide mapping of short-lived proteins in human cells. Molecular cell, 81, 4722–4735 e4725.

68 Cui, Q., Liu, Z. and Bai, G. (2024) Friend or foe: The role of stress granule in neurodegenerative disease. Neuron, 112, 2464–2485.

69 Anderson, P. and Kedersha, N. (2002) Visibly stressed: the role of eIF2, TIA-1, and stress granules in protein translation. Cell Stress Chaperones, 7, 213–221.

70 Youn, J.Y., Dunham, W.H., Hong, S.J., Knight, J.D.R., Bashkurov, M., Chen, G.I., Bagci, H., Rathod, B., MacLeod, G., Eng, S.W.M. et al. (2018) High-Density Proximity Mapping Reveals the Subcellular Organization of mRNA-Associated Granules and Bodies. Molecular cell, 69, 517–532 e511.

71 Markmiller, S., Soltanieh, S., Server, K.L., Mak, R., Jin, W., Fang, M.Y., Luo, E.C., Krach, F., Yang, D., Sen, A. et al. (2018) Context-Dependent and Disease-Specific Diversity in Protein Interactions within Stress Granules. Cell, 172, 590–604 e513.

72 Dubinski, A., Gagne, M., Peyrard, S., Gordon, D., Talbot, K. and Vande Velde, C. (2023) Stress granule assembly in vivo is deficient in the CNS of mutant TDP-43 ALS mice. Human molecular genetics, 32, 319–332.

73 Zhang, X., Wang, F., Hu, Y., Chen, R., Meng, D., Guo, L., Lv, H., Guan, J. and Jia, Y. (2020) In vivo stress granule misprocessing evidenced in a FUS knock-in ALS mouse model. Brain, 143, 1350–1367.

74 Ratti, A., Gumina, V., Lenzi, P., Bossolasco, P., Fulceri, F., Volpe, C., Bardelli, D., Pregnolato, F., Maraschi, A., Fornai, F. et al. (2020) Chronic stress induces formation of stress granules and pathological TDP-43 aggregates in human ALS fibroblasts and iPSC-motoneurons. Neurobiol Dis, 145, 105051.

75 Smith, J. and Bartel, D.P. (2025) The G3BP stress-granule proteins reinforce the integrated stress response translation programme. Nat Cell Biol, 28,135–148.

76 Mercado, G. and Hetz, C. (2017) Drug repurposing to target proteostasis and prevent neurodegeneration: accelerating translational efforts. Brain, 140, 1544–1547.

77 Sidrauski, C., McGeachy, A.M., Ingolia, N.T. and Walter, P. (2015) The small molecule ISRIB reverses the effects of eIF2alpha phosphorylation on translation and stress granule assembly. Elife, 4.

78 Flores, B.N., Yu, S.B., Cohen, I.V., Fanok, M.H., Luan, W., Maciuca, R.D., Sun, L.D., Tsai, R.M., Vissers, M., Smits, L. et al. (2025) Investigational eIF2B activator DNL343 modulates the integrated stress response in preclinical models of TDP-43 pathology and individuals with ALS in a randomized clinical trial. Nat Commun, 16, 7690.

79 Das, I., Krzyzosiak, A., Schneider, K., Wrabetz, L., D’Antonio, M., Barry, N., Sigurdardottir, A. and Bertolotti, A. (2015) Preventing proteostasis diseases by selective inhibition of a phosphatase regulatory subunit. Science, 348, 239–242.

80 Halliday, M., Radford, H., Sekine, Y., Moreno, J., Verity, N., le Quesne, J., Ortori, C.A., Barrett, D.A., Fromont, C., Fischer, P.M., et al. (2015) Partial restoration of protein synthesis rates by the small molecule ISRIB prevents neurodegeneration without pancreatic toxicity. Cell Death Dis, 6, e1672.

81 Bjork, R.T., Mortimore, N.P., Loganathan, S. and Zarnescu, D.C. (2022) Dysregulation of Translation in TDP-43 Proteinopathies: Deficits in the RNA Supply Chain and Local Protein Production. Front Neurosci, 16, 840357.

82 Freibaum, B.D., Chitta, R.K., High, A.A. and Taylor, J.P. (2010) Global analysis of TDP-43 interacting proteins reveals strong association with RNA splicing and translation machinery. Journal of proteome research, 9, 1104–1120.

83 Dominov, J. A., Madigan, L.A., Whitt, J.P., Rademacher, K.L., Webster, K.M., Zhang, H., Banno, H., Tang, S., et al. (2023) Up-regulation of cholesterol synthesis pathways and limited neurodegeneration in a knock-in Sod1 mutant mouse model of ALS. bioRxiv 2023.05.05.539444; doi: 10.1101/2023.05.05.539444

