## Supplementary materials for "STMN2 protein depletion via translation deficits and stress granules and its compensation in ALS"

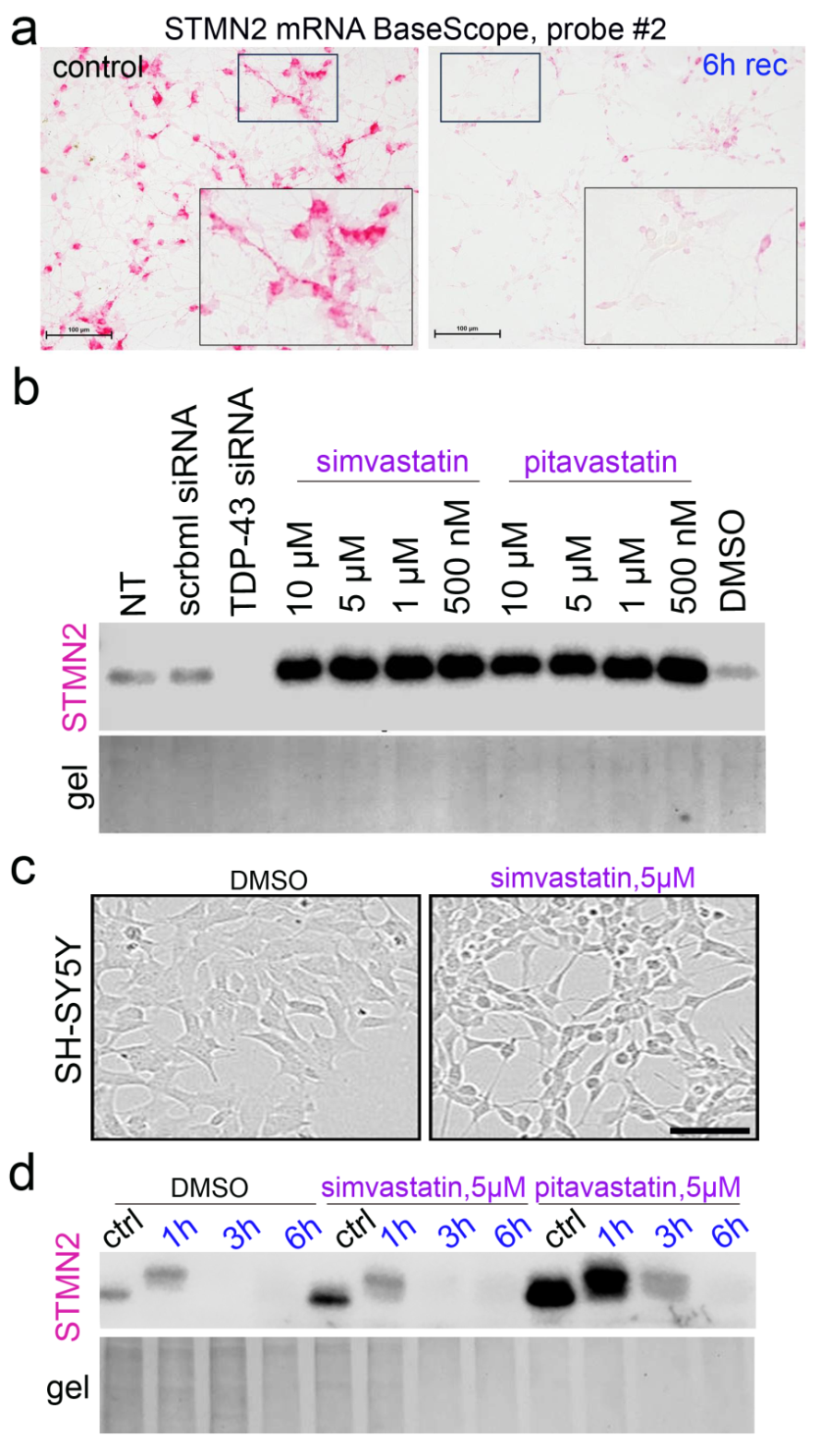

**Figure S1. Characterisation of STMN2 mRNA and protein depletion under acute stress.**

**a,** STMN2 mRNA depletion during the recovery from NaAsO_2_ stress as demonstrated by BaseScope ISH. Probe cat#1048231-C1 was used. Scale bar, 100 µm.

**b,** STMN2 protein is upregulated by statin treatment. TDP-43 siRNA-transfected cells were included as a control. Cells were treated for 16 h. Representative western blot is shown (experiment repeated 3 times).

**c,** High statin concentrations (>500 nM) lead to changes in SH-SY5Y cell morphology, conferring a more “neuronal” phenotype. Representative images are shown. Scale bar, 50 µm.

**d,** STMN2 protein is cleared during stress even when its pre-stress level was elevated by statin pre-treatment. Cells were pre-treated with the indicated concentrations of statins for 16 h, before being subjected to arsenite stress and recovery. Representative western blot is shown (experiment repeated 3 times).

SH-SY5Y cells were used in these studies.

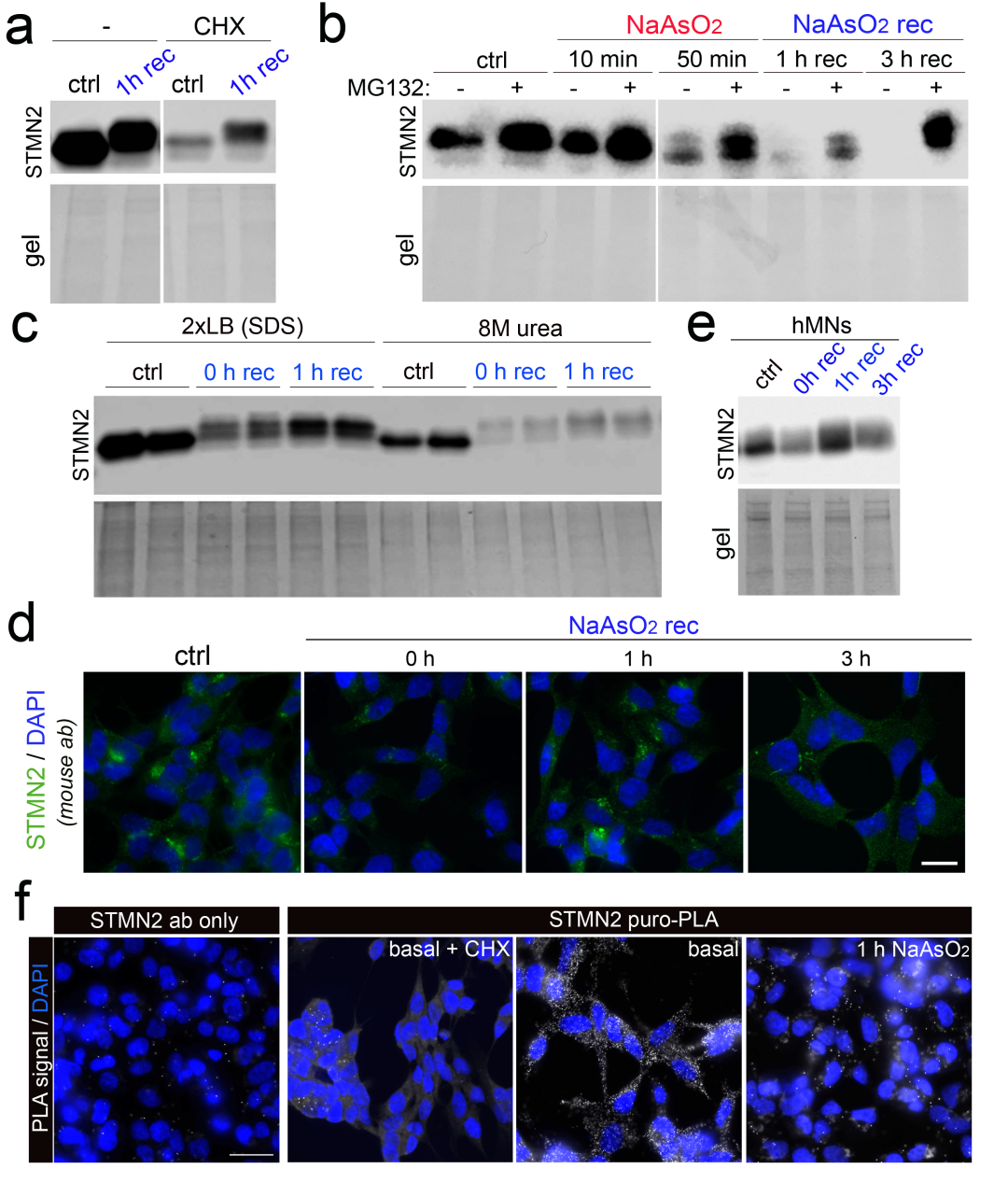

**Figure S2. Characterisation of the dynamic STMN2 protein regulation early in stress.**

**a,** Translation block with CHX leads to STMN2 depletion, however its clearance slows down after 1 h of recovery from arsenite, indicative of declining protein degradation. Cells were pre-treated with CHX for 2 h before arsenite stress, with continued presence of CHX in the media during the recovery. Representative western blot is shown.

**b,** Proteasome inhibition prevents STMN2 protein clearance both early and late in stress. Cells were pre-treated with MG132 for 1 h prior to arsenite addition, with continued presence of MG132 in the media during the recovery. Representative western blot is shown.

**c,** Fluctuations of STMN2 protein level during the recovery from stress are not due to changes in its solubility. Urea-containing buffer was used to solubilise samples prior to western blot. Note that STMN2 protein levels are still higher at 1 h recovery compared to 0 h recovery (=1 h of stress) in the sample set prepared with 8M urea. Representative image is shown (experiment repeated 3 times).

**d,** STMN2 protein level is transiently restored at 1 h of recovery from arsenite stress. A mouse monoclonal antibody for STMN2 was used. Representative images are shown. Scale bar, 10 µm.

**e**, STMN2 protein level is transiently restored during the recovery from arsenite stress in human motor neurons. A mouse monoclonal antibody for STMN2 was used. Representative blot is shown, N=3.

**f,** Puro-PLA can be used to detect and measure STMN2 translation. “STMN2 ab only” – puromycin antibody was excluded during staining, leading to no specific signal. CHX was used as an additional negative control. Note attenuated translation after 1 h of arsenite treatment. Scale bar, 20 µm.

SH-SY5Y cells were used in these studies, except in *e*.

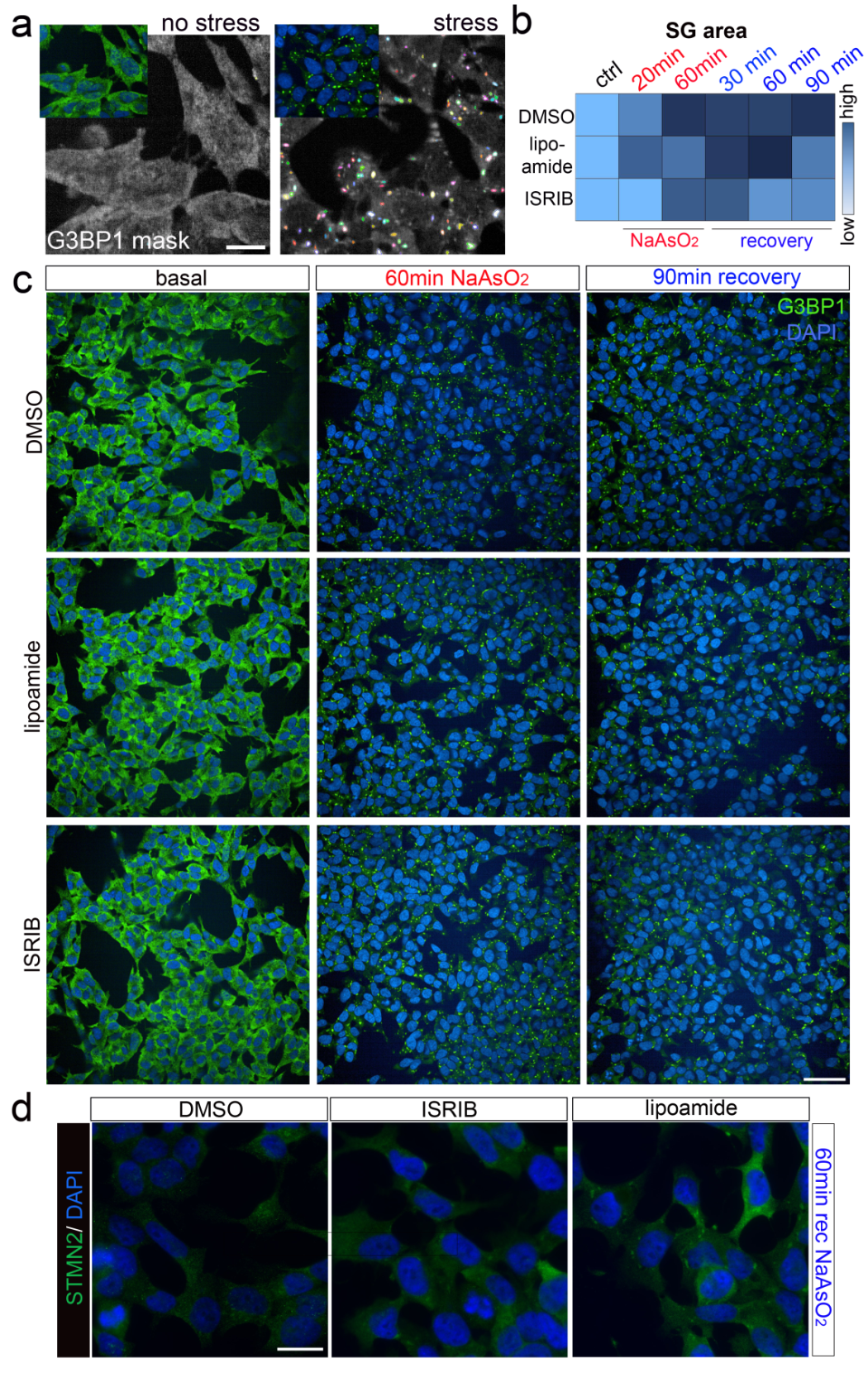

**Figure S3. Modulation of SG assembly affects STMN2 levels during stress.**

**a,** SG analysis on Opera Phenix/Harmony. Representative SG mask is shown. Scale bar, 20 µm.

**b,c,** SG analysis in cells pre-treated with ISRIB or lipoamide during the recovery from NaAsO_2_. An example of a heatmap of the SG area per well for all conditions (b) and representative images for select conditions (c) are shown. Scale bar, 50 µm.

**d,** ISRIB or lipoamide pre-treatment partially prevents STMN2 protein clearance during stress, as detected by immunocytochemistry. Representative images are shown. Scale bar, 10 µm.

SH-SY5Y cells were used in these studies.

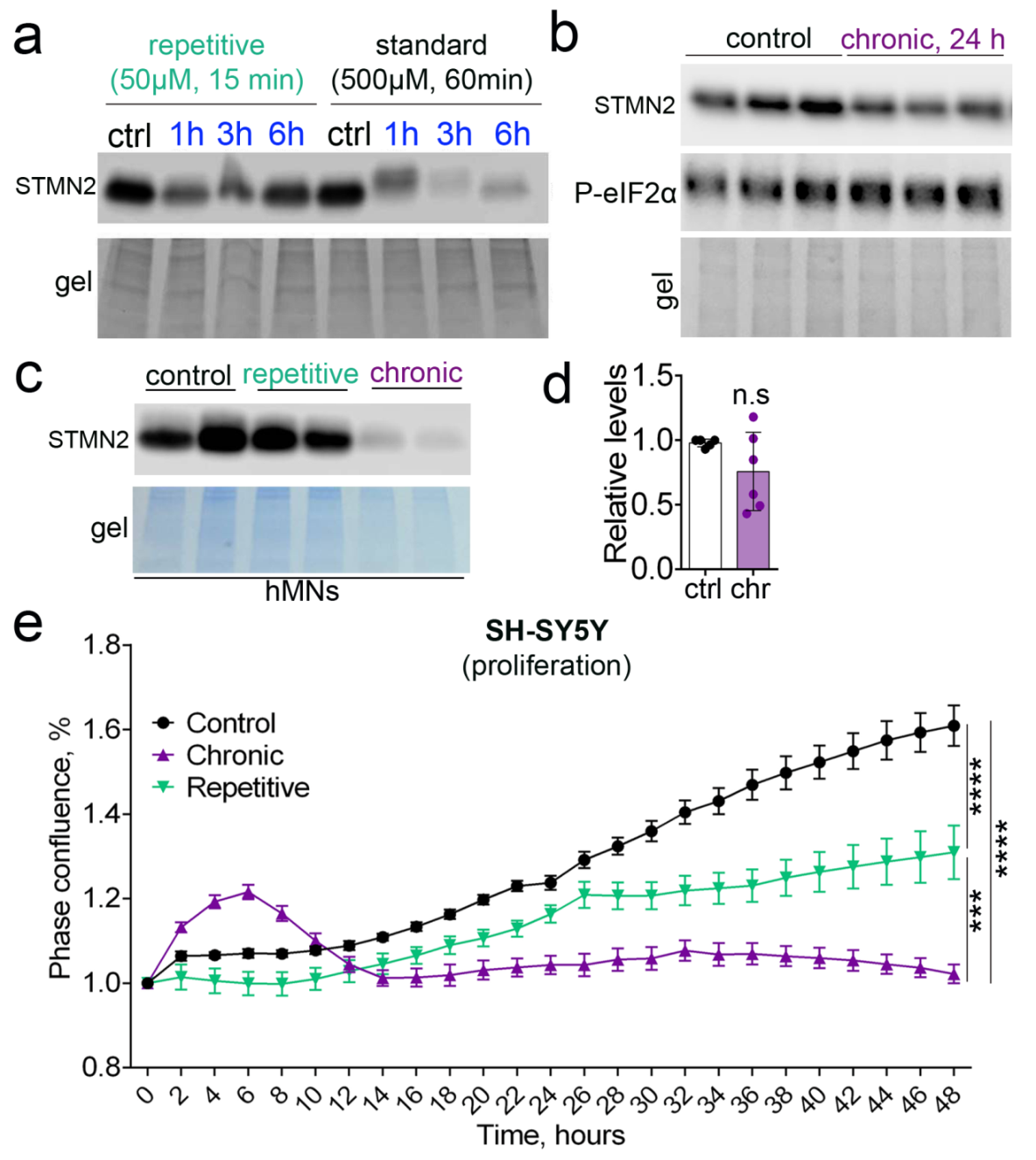

**Figure S4. STMN2 regulation under different stress paradigms.**

**a,** Characterisation of the repetitive stress paradigm. Note that STMN2 protein shows depletion in response to a mild acute stress used in this protocol, albeit less dramatically than in the standard acute stress (single stress) paradigm. Representative western blot is shown.

**b,** Chronic stress reduces STMN2 protein level at the 24 h time-point and promotes eIF2α phosphorylation. Representative western blot is shown.

**c,** Chronic but not repetitive stress reduces STMN2 protein level in human motor neurons. Representative western blot is shown.

**d,** Chronic (chr) stress does not significantly reduce STMN2 mRNA, as analysed by qRT-PCR. N=6. n.s., non-significant.

**e,** Chronic and repetitive stresses both suppress cellular proliferation, with chronic stress being significantly more detrimental. N=4, ***p<0.001, ****p<0.0001, two-way ANOVA with Tukey post-hoc test.

SH-SY5Y cells were used.

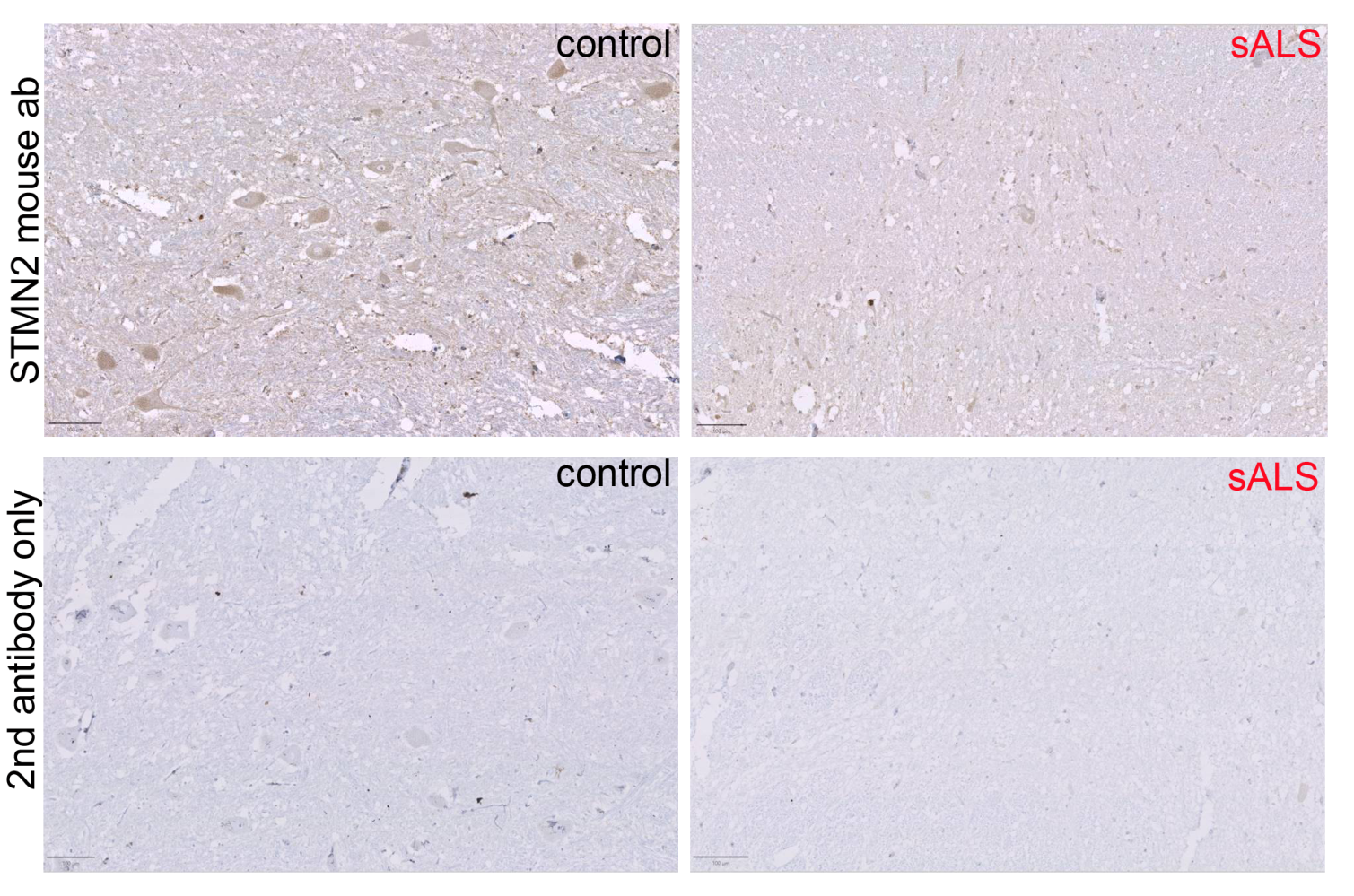

**Figure S5. STMN2 antibody validation for immunohistochemistry in human tissue.**

Spinal cord sections were used. Representative images are shown. Scale bars, 100 μm.

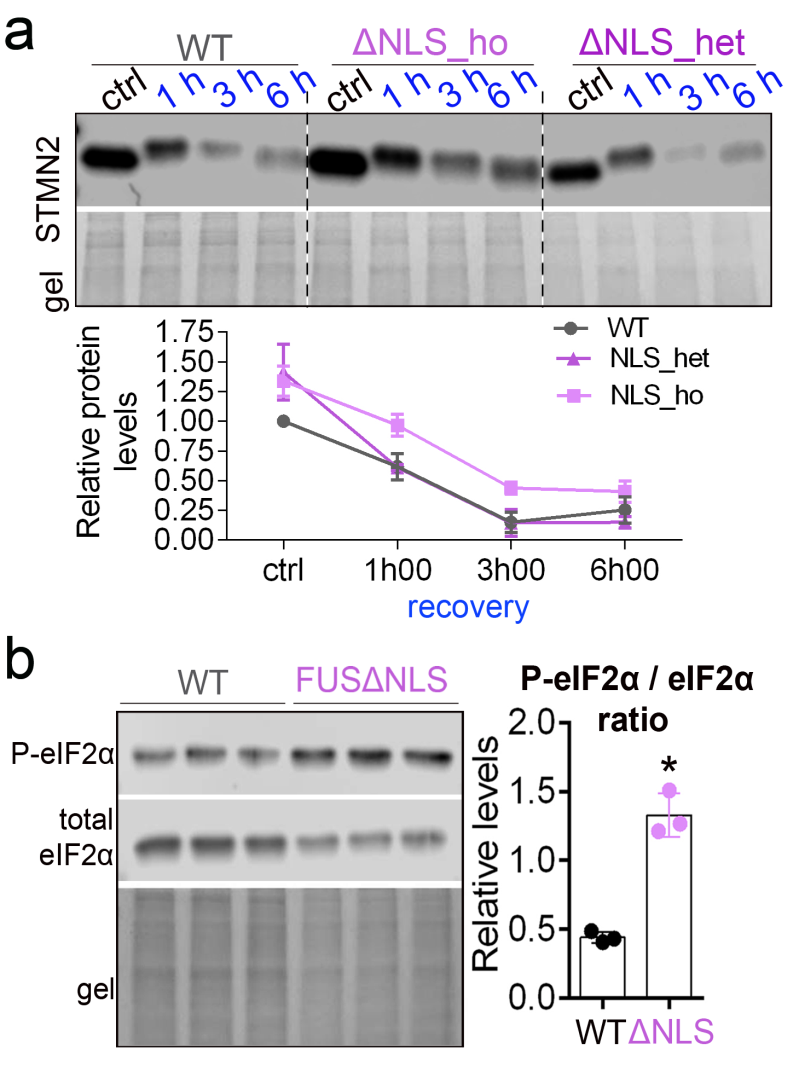

**Figure S6. STMN2 regulation in ALS-FUS cell models.**

**a,** STMN2 protein levels during stress response in ALS-FUS lines.

**b,** ALS-FUS lines are characterised by activated stress response, as demonstrated by elevated phosphor-eIF2α levels. A homozygous line was used. *p<0.05, Mann-Whitney *U* test.

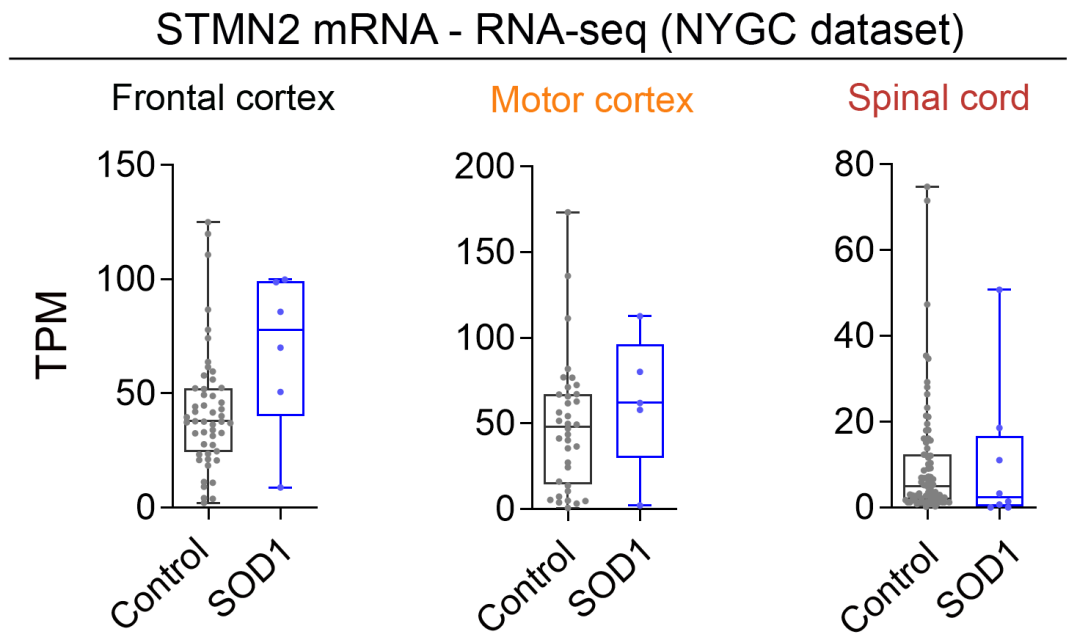

**Figure S7. STMN2 mRNA analysis in ALS-SOD1 post-mortem tissue.**

NYGC ALS consortium bulk RNA-seq data was used.

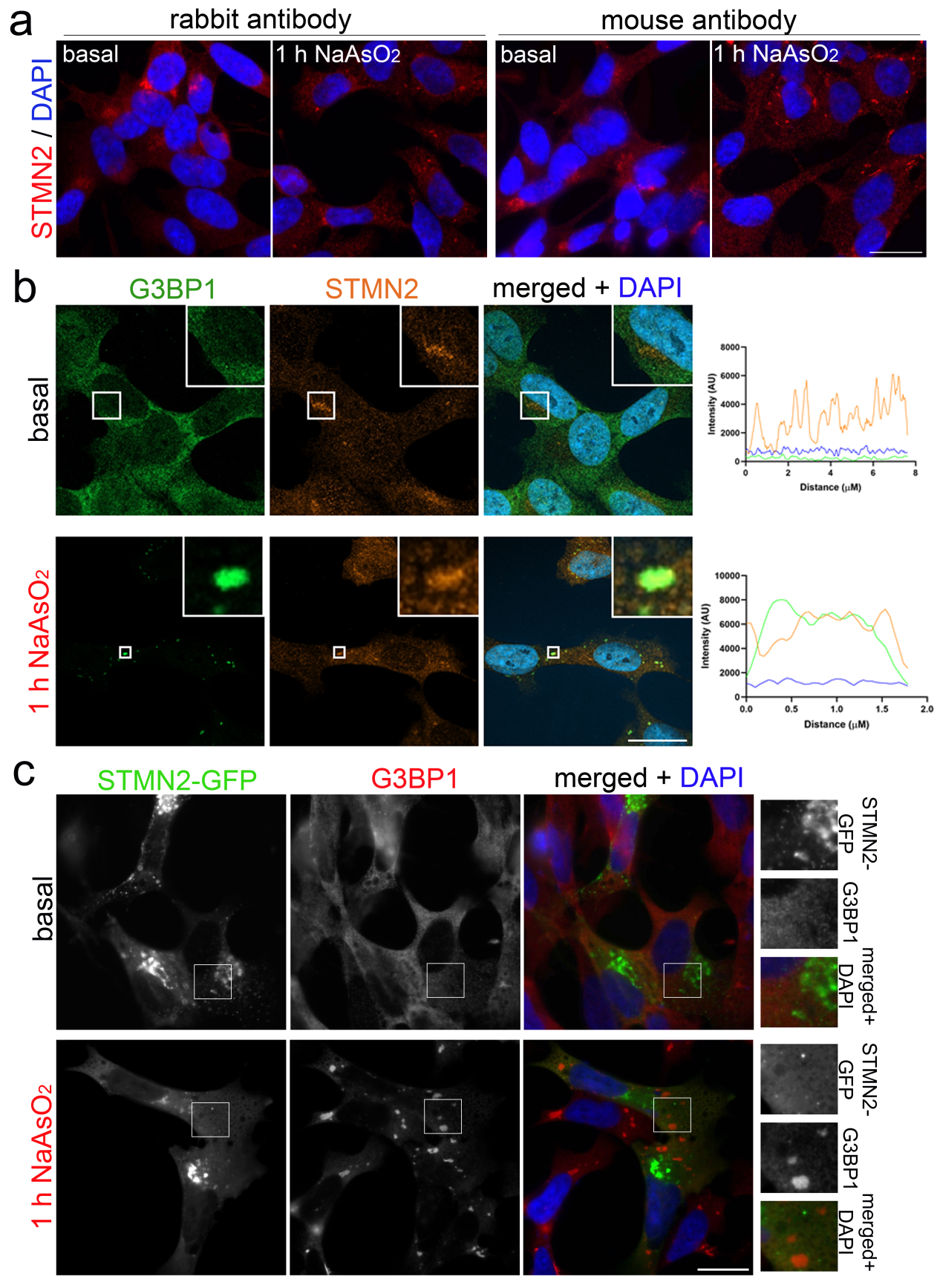

**Figure S8. Analysis of STMN2 protein localisation to stress granules.**

**a,** STMN2 protein is enriched in cytoplasmic foci in stressed cells. Cells were stressed with NaAsO_2_ and analysed during the recovery, using two antibodies.

**b,** STMN2-positive cytoplasmic foci are G3BP1-positive.

**c,** However, transiently expressed STMN2-GFP is not enriched in SGs. Cells were analysed 24 h post-transfection.

SH-SY5Y cells were used in these studies. Representative images are shown. Scale bars, 10 μm.

**Table S1. Sequences of RNA-FISH probes for STMN2 mRNA**

| Probe # | Sequence | Position | %GC |
| --- | --- | --- | --- |
| 1 | ccattgctgttttagcca | 2 | 44.00% |
| 2 | ccttcattttttccttgt | 23 | 33.00% |
| 3 | tcagtgacagcatggaca | 44 | 50.00% |
| 4 | ccgggtaaaagcaagagc | 65 | 56.00% |
| 5 | gatgttgatgttgcgagg | 85 | 50.00% |
| 6 | tcacttccatatcatcgt | 110 | 39.00% |
| 7 | gaggcacgtttgttgatt | 132 | 44.00% |
| 8 | tcagctcaaaagcctggc | 152 | 56.00% |
| 9 | taggagatggtggcttca | 173 | 50.00% |
| 10 | aagttcgtggggcttctg | 194 | 56.00% |
| 11 | gtctttcttctttggaga | 217 | 39.00% |
| 12 | tggatctcctccagggac | 237 | 61.00% |
| 13 | tctgcagcctccagtttc | 258 | 56.00% |
| 14 | cctcctgagactttcttc | 281 | 50.00% |
| 15 | ccaattgtttcagcacct | 302 | 44.00% |
| 16 | ctcgtgttccctcttctc | 322 | 56.00% |
| 17 | gccttctgaaggacttct | 342 | 50.00% |
| 18 | aagttgttgttctcctcc | 363 | 44.00% |
| 19 | ttttcctccgccatcttg | 384 | 50.00% |
| 20 | gttccattttcaggatca | 404 | 39.00% |
| 21 | gctagattagcctcacgg | 435 | 56.00% |
| 22 | cttttcctgcagacgttc | 463 | 50.00% |
| 23 | gagttccttgttcctgcg | 502 | 56.00% |
| 24 | cagccagacagttcaacc | 522 | 56.00% |
